# Real-time brain-state-coupled cortico-cortical paired associative stimulation of cognitive networks

**DOI:** 10.64898/2026.05.01.722353

**Authors:** D. Blair Jovellar, Sonia Turrini, Paolo Belardinelli, Olivier Roy, Emiliano Santarnecchi, Ulf Ziemann

## Abstract

Brain networks coordinate distributed neuronal assemblies to support cognition. Spike-timing-dependent plasticity (STDP) and neuronal oscillations are key substrates for state-gated learning rules that shape network coupling and cognitive operations; nonetheless, how STDP mechanisms interact with neuronal oscillations is largely unexplored in humans. Cortico-cortical paired associative stimulation (ccPAS) provides a non-invasive system-level model of associative timing rules by pairing dual-site transcranial magnetic stimulation (TMS) across axonally connected regions with an inter-stimulus interval matched to pathway conduction. Here we: 1) synthesize ccPAS applications and barriers to brain-state-coupled implementation in cognitive networks; 2) provide an actionable roadmap for real-time state estimation, targeting, and dual-site parameter selection; and 3) demonstrate a novel implementation of theta-phase-locked fronto-parietal (FP) ccPAS with concurrent EEG in adult human participants.

We tested whether ccPAS delivered at the positive phase of ongoing theta (POS) induces distinct changes in evoked EEG activity and FP connectivity compared to phase-uncoupled ccPAS (RAND) and phase-locked single-site prefrontal (PREF) controls. At the evoked level, POS produced a fronto-central polarity reversal of the canonical N45 component and a right parieto-temporal negativity relative to both controls. At the network level, POS induced frequency-specific reconfigurations in post-intervention connectivity beyond either control ingredient alone. Together, these changes in evoked activity and rapid network reconfiguration provide the first empirical evidence consistent with phase-gated STDP in humans—whereby oscillatory phase gates cortical excitability and modulates STDP efficacy—emerging as short-term network-level expression. Future work will assess long-term plasticity by tracking connectivity at later time points and testing for concomitant behavioral effects.

**Significance:** The real-time brain state critically shapes how plasticity mechanisms are expressed in response to brain stimulation. This article provides a forward-looking synthesis of the scientific and technical challenges associated with ccPAS—an STDP induction model in the human cortex—and outlines the steps required to advance it toward real-time brain-state-coupled implementation. To our knowledge, this is the first application of brain-state-coupled ccPAS within a cognitive network. By personalizing stimulation to the individual’s ongoing neural state, this approach may reduce variability, limit off-target effects, and enhance plasticity induction. Ultimately—by modulating network-level function in a brain-state-dependent manner—this technique could augment therapeutic outcomes in disorders marked by network dysfunction such as ADHD, Alzheimer’s disease, and major depressive disorder, potentially maximizing efficacy in patients unresponsive to existing treatments.

## Introduction

Many cognitive disorders—such as Alzheimer’s disease, ADHD, and major depression—are characterized by network-level dysfunction rather than deficits confined to isolated brain regions (Gao et al., 2019; Palop and Mucke, 2016; Kaiser et al., 2015). Brain stimulation is a promising tool for modulating large-scale networks and improving cognitive performance. Indeed, recent advances in non-invasive brain stimulation (NIBS) allow not only for the modulation of activity within individual brain regions, but also for the manipulation of connectivity between them via Hebbian plasticity (Pirazzini et al., 2026; Trajkovic et al., 2023; Momi et al., 2022). Cortico-cortical paired associative stimulation (ccPAS) uses dual-site TMS to stimulate superficial cortical regions with the aim of modulating synaptic efficacy along targeted axonal pathways (Di Luzio et al., 2024; Hernandez-Pavon et al., 2022). While ccPAS is applicable across motor, sensory, or cognitive domains, the present work centers on its implementation in cognitive networks—such as fronto-parietal, cingulo-opercular, and default mode systems—which are large-scale association networks supporting higher-order, transmodal functions including working memory, attention, decision-making, and cognitive control. These networks are distinguished from unimodal sensory or motor systems by their long-range connectivity, integrative computational roles, and capacity for flexible reconfiguration across task demands (Mesulam, 1998; Power et al., 2011; Herbet & Duffau, 2020; Margulies et al., 2016). This functional distinction motivates the development of brain-state-dependent ccPAS protocols capable of selectively modulating connectivity within association networks, where the timing and configuration of large-scale dynamics critically shape cognitive operations.

ccPAS provides a network-level implementation based on STDP, a Hebbian mechanism wherein the precise timing of pre- and post-synaptic activity strongly biases the induction of long-term potentiation (LTP) or long-term depression (LTD) (Markram et al., 1997, 2012; Bi and Poo, 1998; Feldman, 2000; Nishiyama et al., 2000; Sjöström et al., 2001; Wittenberg and Wang, 2006). In ccPAS, each TMS pulse evokes a synchronized population volley, and sequential stimulation of two interconnected regions produces pre- and post-synaptic volleys that travel along shared pathways and converge with millisecond precision. A well-chosen inter-stimulus interval (ISI) allows these volleys to arrive within an STDP-relevant temporal window, thereby modifying synaptic efficacy across the many synapses forming the targeted cortico-cortical pathway. Notably, mechanistic studies show that spike timing alone does not fully determine the direction or durability of plasticity: pairing structure, repetition frequency, and total number of pairings jointly shape whether LTP, LTD, or no change is expressed (Wittenberg & Wang, 2006; Cui et al., 2018; Anisimova et al., 2022). Consistent with established STDP physiology, resulting plasticity may be expressed at pre-synaptic, post-synaptic, or combined loci, mediated by changes in release probability, vesicle dynamics, AMPAR trafficking and retrograde signaling (Bi & Poo, 1998; Markram et al., 1997; Sjöström et al., 2001; Kessels & Malinow, 2009; Fino et al., 2008; Lüscher & Malenka, 2012). While previous ccPAS studies (Arai et al., 2011; Lu et al., 2012; Chao et al., 2013) reported effects mainly in the second stimulated region (i.e., M1), this likely reflects the specificity of the outcome measure: motor evoked potentials (MEPs) are sensitive only to the excitability of corticospinal neurons specifically in M1 that project to spinal motor neurons—not to activity changes in upstream presynaptic sites. In contrast, studies using TMS-Evoked Potentials (TEPs), which more directly assay cortical reactivity, have shown that ccPAS can modulate activity in both stimulated regions (Casula et al., 2016; Veniero et al., 2013), supporting the view that ccPAS induces pathway-specific plasticity distributed across the targeted circuit.

ccPAS has shown efficacy in modulating a range of higher-order cognitive functions. Acting on connectivity within the FP executive network can modify decision making (Nord et al., 2019) and fluid intelligence (Momi et al., 2020), while modulating bilateral lateral prefrontal connectivity can regulate emotional reactivity (Zibman et al., 2019). While promising, all previous ccPAS studies have disregarded the ongoing brain state during stimulation—which is a critical modulator of NIBS outcomes (Zrenner et al., 2018; Romei et al., 2016). Optimizing ccPAS for cognitive networks therefore requires understanding how to align stimulation with endogenous states that facilitate plasticity.

Applying ccPAS across two cortical regions increases methodological complexity and expands the parameter space for successful implementation. Cognitive network applications pose unique challenges not encountered in motor systems, such as the absence of a real-time physiological readout analogous to MEPs. Brain-state-coupled ccPAS introduces further considerations regarding real-time monitoring, detection, and synchronization with endogenous activity.

In this paper, we *(i)* review the state-of-the-art and current challenges in brain-state-coupled ccPAS, focusing on target selection and stimulation parameters; *(ii)* provide a practical roadmap for implementing brain-state-coupled ccPAS; and *(iii)* demonstrate its feasibility in an FP network supporting working memory. We tested whether ccPAS delivered at the positive phase of ongoing theta (POS) induces distinct changes in evoked EEG activity and FP connectivity compared to phase-uncoupled ccPAS (RAND) and phase-locked single-site prefrontal (PREF) controls. The POS ccPAS-induced changes in evoked activity and rapid network reconfiguration provide the first empirical evidence consistent with phase-gated STDP in humans—whereby oscillatory phase gates cortical excitability and modulates STDP efficacy—emerging as short-term network-level expression. By articulating strategies for brain-state-coupled ccPAS application, we aim to provide a practical and conceptual framework for researchers seeking to implement this complex yet powerful technique for modulating brain networks.

## I. Brain-state-coupled ccPAS: State-of-the-art and current challenges

Implementation of brain-state-coupled ccPAS requires scrutinizing and refining experimental approaches which include considerations for: *i)* target optimization, *ii)* stimulus parameter characterization, and *iii)* ccPAS TMS-related artifact mitigation

### Target considerations

Defining the correct targets is critical to the success of brain-state-coupled ccPAS. Here, target refers to two distinct components: the anatomical target, i.e., selecting the correct stimulation nodes (regions) in the network of interest, and the target brain state, i.e., determining an appropriate neurophysiological brain state marker to trigger the paired stimulation.

#### Anatomical target

The aim of ccPAS is to manipulate connectivity between network nodes; thus, it is necessary to define two targets whose connectivity, if enhanced or decreased, would lead to specific measurable neurophysiological or behavioral aftereffects. Selecting what nodes within a network to stimulate is not trivial, and the choice can be informed by multiple strategies. One option would be to target areas belonging to specific cognitive networks whose enhanced co-activation correlates with increased performance in selected tasks, based on previous literature. For instance, this is the approach adopted by multiple studies that have targeted FP nodes (Zibman et al., 2019; Nord et al., 2019). This method has some indubitable advantages, such as defining common and shared stimulation sites for all individuals recruited in a study, which greatly simplifies the interpretation of results. However, it brushes over individual differences in the cortical correlates of cognition, which are known to be highly impactful, as functional connectivity patterns are so peculiar to each individual that they can act as individual “fingerprints” (Nentwich et al., 2020; Finn et al., 2015).

A second targeting approach would be to select paired stimulation sites based on individual functional connectivity maps whereby regions that are activated during cognitive tasks are engaged using TMS (for a review see Fox et al., 2012). This method of target selection has been applied to ccPAS in the cognitive domain by two studies (Momi et al., 2020; Santarnecchi et al., 2018), which chose targets based on fMRI connectivity data collected at baseline, and obtained significant results on behavior, undirected connectivity, and effective connectivity. Still, this selection method has its setbacks. By design, this leads to varying target sites between participants (Momi et al., 2020), which could make the setting of shared stimulation parameters (see below) and the interpretation of results more challenging. Additionally, functional connectivity is significantly different when measured with fMRI as opposed to EEG (Nentwich et al., 2020) due to the inherent differences in spatial and temporal resolution; therefore, the definition of ideal targets would vary based on the imaging technique used to extract functional connectivity maps.

A third approach is anatomically guided TMS targeting, which incorporates information from diffusion MRI and/or structural MRI. Diffusion MRI enables the identification of structurally connected brain regions via tractography, while structural MRI provides high-resolution cortical morphology to optimize coil placement.

Target selection based on diffusion MRI can be performed using offline tractography (Momi et al., 2021) or emerging real-time techniques (Aydogan et al., 2025). Evidence indicates that TMS-induced activation preferentially propagates along white matter pathways (Momi et al., 2021), and that the structural connectome strongly constrains large-scale patterns of brain coactivation (Horn et al., 2014; Honey et al., 2009; Skudlarski et al., 2008). These relationships are observed across both task-based and resting-state functional connectivity (Hermundstad et al., 2013), suggesting that structurally informed targeting may implicitly reflect functional network architecture.

In parallel, structural MRI supports targeting of cortical regions with more favorable biophysical properties. Specifically, it allows for adjustment based on scalp-to-cortex distance (Stokes et al., 2005) and facilitates precise localization of gyral crowns, where the induced electric field is stronger and more orthogonally aligned with cortical columns—compared to sulcal walls, where stimulation is typically less effective (Klomjai et al., 2015).

#### Brain state target

The second critical component of brain-state-coupled ccPAS is identifying a robust brain state marker. We define a brain state as a momentary operating condition of the brain, reflecting the current configuration of neural activity and the physiological and behavioral context in which information processing occurs. Since brain stimulation applied during different states yields distinct effects on brain activity (Bradley et al., 2022), a reliable brain state marker is necessary to detect when the desired state is present and to precisely couple stimulation to that state.

Broadly, brain states can be characterized across four overlapping domains:

- **Behavioral states** refer to externally observable conditions such as wakefulness, sleep, or engagement in a sensory task (Grimm et al., 2024; Tagliazucchi & Laufs, 2014).
- **Cognitive and affective states** reflect internal processes like attention, perception, or mood—examples include focused vs. distracted attention (Christoff et al., 2009), depressive states (Lynch et a., 2024), and changes in affective arousal (Zhang et al., 2025).
- **Neurophysiological states** capture patterns of brain activity across multiple spatial and temporal scales—typically assessed using electrophysiological methods such as EEG and MEG, or hemodynamic techniques like fMRI. While EEG and MEG provide direct, high-temporal-resolution measures of oscillatory rhythms and cortical excitability, fMRI is widely used to infer neurophysiological processes through analyses of functional connectivity, task-evoked activation, and dynamic network reconfiguration (Klimesch, 1999; Fox et al., 2005; Silvanto et al., 2008).
- **Biochemical or neurochemical states** reflect the molecular, metabolic, and vascular environment of the brain. These states are typically quantified through direct molecular imaging techniques (e.g., PET, MRS) or indirect proxies that reflect neurochemical and metabolic activity. Examples include neurotransmitter concentrations (such as dopamine, GABA, and glutamate), cerebral oxygenation and hemodynamic responses measured via BOLD fMRI or fNIRS, and other markers of energy metabolism and neurovascular coupling (Grace, 2016; Stagg et al., 2011; Cui et al., 2011). While not direct measures of neuronal signaling, these indices provide critical insight into the biochemical and vascular substrates that modulate brain function and may serve as state indicators under certain conditions.

Because of this conceptual plurality, the choice of brain state markers must be closely aligned with both the goals and constraints of the experimental paradigm. In the context of brain-state-dependent neuromodulation, neurophysiological states are typically targeted using EEG. EEG is favored for its *(i)* high temporal resolution, *(ii)* compatibility with real-time monitoring, *(iii)* non-invasiveness, *(iv)* cost-efficiency, and, crucially, *(v)* ability to capture transient excitability-state fluctuations relevant to stimulation efficacy (Thut et al., 2017; Sun et al., 2024).

Within this framework, we focus on neural oscillations—rhythmic fluctuations in population-level excitability that coordinate brain activity across time and space (Buzsáki & Draguhn, 2004). Oscillatory dynamics offer a continuous and temporally specific signal from which key brain states can be derived. EEG features such as phase-locked activity, within- and cross-frequency coupling, and graph-theoretic network metrics (e.g., centrality, degree, path length) provide a rich window into large-scale, spatiotemporal dynamics at behaviorally relevant timescales.

Brain states during cognitive functions have been well-characterized using phase, connectivity, and network measures. A critical feature of the applicability of such measures for brain state-dependent stimulation is whether they can be instantaneously calculated (within hundreds of milliseconds to ∼1 second). In this regard, oscillatory phase as a brain state marker for state-dependent stimulation stands out as it can be easily and quickly calculated in real-time. Phase-locked stimulation has been applied in conventional single site TMS (Hassan et al., 2023; Zrenner et al., 2018), transcranial electrical stimulation (Zarubin et al., 2020), and even optogenetic stimulation in animals (Siegle et al., 2014). Other neurophysiological measures that have been calculated in real-time are phase-amplitude coupling (i.e., local mutual information) (Martinez-Cancino et al., 2019) and phase synchronization (i.e., phase locking value) (Pieramico et al., 2023). Single-trial phase synchronization has also recently been evaluated and targeted using real-time EEG-TMS in the motor system (Vetter et al., 2023). Because phase- locked stimulation is well-characterized, we use it in a novel brain-state-dependent ccPAS protocol (Section 3).

### Stimulus parameters

Exogenously inducing STDP in selected cortical paths is far from trivial—a multitude of technical and practical considerations come into play. Assuming the target nodes and brain states have been defined, it is then necessary to define how to repeatedly and coherently stimulate them to modify their connectivity. The parameters to set (Figure 1) are multiple: 1) the stimulation order (i.e., which brain region is stimulated first versus second); 2) the inter-stimulus interval (ISI) between the pulse delivered over the first and second node, 3) the stimulation intensity of both cortical areas, 4) the frequency of paired-pulse delivery (inter-trial interval), and 5) total number of paired pulses.

**Figure 1.**
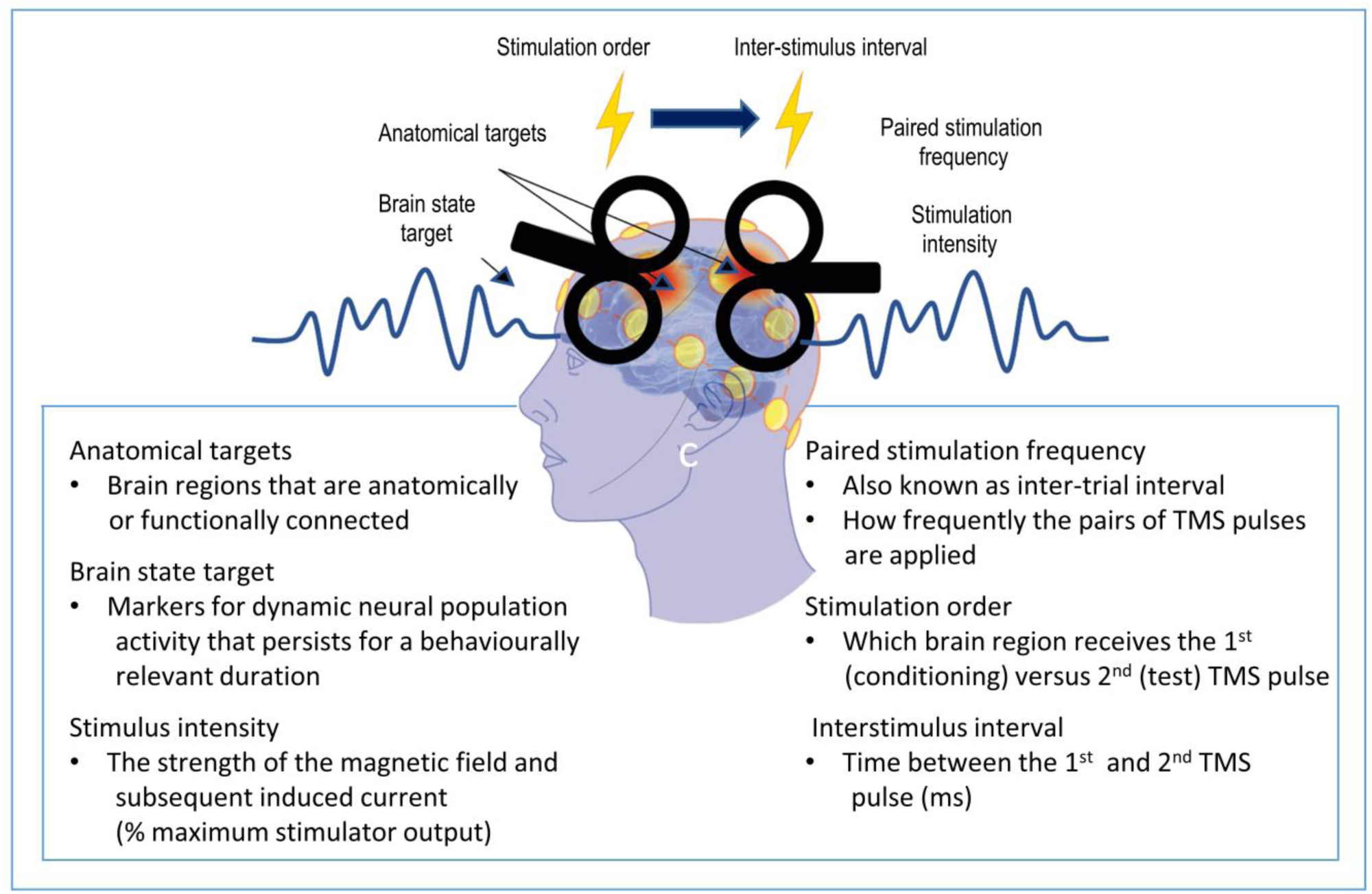
Stimulus parameters for brain- state-coupled cortical paired associative stimulation. Schematic of stimulus parameters with descriptions.

#### Stimulation order

Deciding which brain region to apply the conditioning (first) and test (second) pulse is the most critical parameter to define for ccPAS. As a cortical system-level model of synaptic STDP, the order of stimulation is one of the key determinants of plasticity direction (Markram et al., 1997; 2012). Accordingly, repeatedly stimulating the “pre-synaptic area” first leads to LTP, while stimulating the “post-synaptic area” first results in LTD. How then can we identify the pre-synaptic versus post-synaptic regions for ccPAS in the human cortex? This is a highly challenging question to answer since several interconnected brain regions have reciprocal connections (Diao et al., 2018; Massé et al., 2016; Giarrocco and Averbeck, 2021; Rosenberg et al., 2009). As such, majority of ccPAS studies in various cognitive domains have performed pilot or exploratory experiments first where the paired pulses were applied in both directions across different sessions (Sallie et al., 2023; Kohl et al., 2019; Momi et al., 2020; Nord et al., 2019; Zibman et al., 2019; Santarnecchi et al., 2018; Casula et al., 2016). While performing an initial experiment stimulating in either direction unequivocally shows how ccPAS in each direction affects outcomes (e.g., behavioral or neurophysiologic measures), other strategies can also be used to ascertain the stimulation order. Previous anatomical tract tracing studies in non-human primates and even human tissues (Xia et al., 2024; Mavrovounis et al., 2024; Sivukhina and Jirikowski, 2021; Lanciego and Wouterloud, 2020; Heilingoetter and Jensen, 2016; Sparks et al., 2000) can inform origin (pre) and termination (post) sites. Performing subject-specific white matter tractography (Schiavi et al., 2020; Aydogan et al., 2018; Wang et al., 2015) can also help define pre versus post areas and further refine the coordinates of both stimulation sites in a personalized manner. In the presence of reciprocal connections, white matter imaging studies (Tanner et al., 2023; Góngora et al., 2016) can also reveal any asymmetries (i.e., which direction is more dominant)—noting that any differences in connectivity and integrity may evolve across the lifespan (Cox et al., 2016; Liu et al., 2017; Alkonyi et al., 2011) and correlate with cognitive function (Sha et al., 2023; d’Arbeloff et al., 2019; Mito et al., 2018; Mamah et al., 2010). Moreover, incorporating functional information can help resolve the directionality question, particularly when brain regions have bidirectional connections. One can examine the direction of information flow during a cognitive task that activates both target areas based on functional imaging studies using electrocorticography (ECoG), magnetoencephalography (MEG), EEG, or fMRI (Zhou et al., 2023; Dimakopoulos et al., 2023; Masson and Isik, 2023; Goddard et al., 2022; Schoffelen et al., 2017; Thatcher et al., 2016). Integrating both structural and functional information more effectively informs the stimulation order to elicit desired ccPAS outcomes.

#### Inter-stimulus interval

The precise timing of stimulation of the two cortical targets is the second most important stimulus parameter to define in ccPAS applications. Indeed, the strength of the STDP effect depends on the correct selection of the ISI between the two pulses and, when sub-optimal, STDP effects can be reduced (Chiappini et al., 2022) or nullified (Lazari et al., 2022; Nord et al., 2019; Casula et al., 2016; Johnen et al., 2015; Koganemaru et al., 2009). ccPAS studies in the motor system (Turrini et al., 2023 a,b, 2022; Koch et al., 2020, 2013; Fiori et al., 2018; Johnen et al., 2015; Veniero et al., 2013) have mostly been informed by dual coil effective connectivity studies, that measure MEP amplitude modulation as a direct readout, to define ISIs. Lacking a direct readout such as the MEP, ccPAS studies targeting the FP network, thus far, have derived their ISIs from work conducted in the motor system based on the assumption that the FP regions being stimulated have similar length oligosynaptic connections as the motor targets of previous studies (Momi et al., 2020; Nord et al., 2019, Casula et al., 2016). More recently, one study adopted the combination of TMS and EEG, delivering TMS over one cortical area and recording evoked potentials from a remote site to characterize the cortico-cortical conduction timing between the two, and tailored a subsequent ccPAS protocol to the timing of the evoked potentials which resulted in remarkable electrophysiological and behavioral effects (Borgomaneri et al., 2023). Another study defined the ISI based on the communication-through-coherence framework. By targeting the pathway connecting the left and right V5 areas, which oscillates in the gamma band (40 Hz), the ISI was calibrated to align with the time lag between the peaks of the gamma oscillatory rhythm (Chiappini et al., 2022). These methods could also be applied to FP and other networks. Alternatively, assuming the selected targets are directly connected by white matter pathways, ISIs could be informed by tractography-based estimates of conduction delay derived from tract length and microstructural properties of the white matter fiber bundle (Caminiti et al., 2013; Mancini et al., 2021).

#### Stimulation intensity

Dual coil TMS studies on effective connectivity conducted mostly in the motor system indicate that different stimulation intensities of a first area exert starkly dissimilar conditioning effects over anatomically connected nodes. In the widely studied premotor-motor network, the conditioning effect exerted by the premotor cortex over the excitability of the primary motor cortex was found to shift from inhibitory to facilitatory based on the intensity of the stimulation delivered over the premotor cortex (Baumer et al., 2009; Civardi et al., 2001). The few ccPAS studies targeting higher cognitive functions have, so far, applied heterogeneous stimulation paradigms, setting the intensities based on individual resting motor thresholds but employing suprathreshold stimulation of both targeted nodes in some cases (Nord et al., 2019; Zibman et al., 2019; Casula et al., 2016) and adopting subthreshold conditioning in others (Momi et al., 2020; Santarnecchi et al., 2018). Basing the stimulus intensity for cognitive regions on motor cortex excitability is inherently a flawed approach due to anatomical and physiological differences (e.g., cytoarchitecture, neurotransmitter receptors, cortex-to-scalp distance, cortical folding structure, etc.) that lead to endogenous variations in excitability between brain regions. To find the optimal intensities for the conditioning and test pulse in cognitive regions, two questions need to be addressed: *i)* what intensity is needed to effectivity activate neurons in each stimulated region (single site TMS intensity), and *ii)* how varying levels of intensities for each region interact. One method for determining the intensity particularly for non-motor regions proposed by Stokes and colleagues (2005) is to scale the motor threshold based on scalp-to-cortex distance, based on a simple mathematical formula. However, this method is likely oversimplified, as it does not take into account other factors, such as cytoarchitecture or individual folding structure.

An emerging approach to TMS intensity dosing is via E-field modeling. This method is particularly relevant for non-motor regions where there is no direct method for evaluating the effects of varying stimulus intensities. An example E-field modeling tool is SimNIBS (Saturnino et al., 2019). While majority of E-field modeling approaches focus on modeling electromagnetic induction that can depolarize cortical neurons in single brain regions (Daneshzand et al., 2021; Xu et al., 2021; Makarov et al., 2020; Stenroos and Koponen, 2019) a method that is pertinent to ccPAS applications embeds structural connectivity based on white matter tractography in the E-field model (Geeter et al., 2016). Geeter and colleagues illustrate the effects of varying stimulation intensities and coil orientations on the spatial distribution of the membrane potentials along fiber tracts and its temporal dynamics. While this method has only been tested using single site stimulation, this approach would be also potentially valuable in ccPAS paradigms, which seek to modulate the activity of interconnected brain regions.

#### Frequency and number of paired pulses

The frequency and number of the delivered paired pulses also play critical roles in the induction of Hebbian plasticity, which requires coherent and repeated stimulation. Previous ccPAS studies in cognitive networks (Casula et al., 2016; Santarnecchi et al., 2018; Nord et al., 2019; Momi et al., 2019; Kohl et al., 2019; Zibman et al., 2019) have applied 100-210 rhythmic paired stimuli with frequencies between 0.1 and 0.25 Hz. More recently, a protocol has used higher total pair count and repetition rate, applying 600 paired pulses at ∼0.33 Hz (Aviram-Friedman et al., 2025). The slower inter-pair frequencies were adopted from early M1 ccPAS studies (Arai et al., 2011; Koch et al., 2013; Veniero et al, 2013) which used low repetition rates to emphasize associative timing while minimizing non-associative rate-based effects (Nord et al., 2019; Santarnecchi et al., 2018). While low frequency ccPAS protocols may be contextualized against single site rTMS—where 1 Hz stimulation has traditionally been associated with LTD-like reduction in cortical excitability (Ziemann et al. 2008), recent evidence showed that 1 Hz and 10 Hz rTMS did not reliably modulate corticospinal excitability (Magnuson et al., 2023). Moreover, 0.2 Hz stimulation was found to increase excitability (Pellicciari et al., 2016), whereas 0.1 Hz did not produce measurable changes even when stimulation was applied for 1 hour (Chen et al., 1997). Accordingly, repetition rate alone is not a reliable predictor of TMS physiological aftereffects, particularly in ccPAS, motivating the need to consider how multiple parameters interact and govern timing-dependent plasticity.

Mechanistic studies demonstrate that pairing structure, repetition rate, and the total number of pairings (Npairings) jointly determine not only the direction of STDP but also its robustness to biological variability (Wittenberg & Wang, 2006; Cui et al., 2018; Anisimova et al., 2022). Potentiation can emerge with relatively few causal pairings—often only tens—when delivered at theta-like frequencies (Wittenberg & Wang, 2006), whereas LTD typically requires a substantially larger volume of activity (∼100–300 pairings) to be expressed reliably (Wittenberg & Wang, 2006; Cui et al., 2018; Anisimova et al., 2022). The persistence of plasticity also depends on frequency and pulse count. Late LTP at CA3-CA1)—which was observable three days after STDP induction— only occurred when the pairing frequency was 5 Hz or when Npairings was increased to ∼250-300, whereas 0.1 Hz pairings with fewer repetitions did not yield durable potentiation (Anisimova et al., 2022). Increasing either the pairing frequency (e.g., 3–5 Hz) or Npairings (≥250–300) also buffers NMDA-mediated LTP against spike-timing jitter (Cui et al., 2018).

Collectively, these studies indicate that LTP and LTD occupy a shared, multidimensional parameter space defined by ISI, frequency, and Npairings. Existing human ccPAS protocols sample only a narrow portion of this biologically effective parameter range, primarily at very low frequencies with modest pair counts. Future protocols may benefit from treating both repetition rate and total pair number as explicitly adjustable parameters for optimizing pathway-specific associative plasticity.

### ccPAS TMS-related artifacts

Paired-pulse protocols like ccPAS introduce unique challenges and amplify existing artifacts compared to single-pulse TMS. This section outlines how these artifacts differ and strategies for mitigating them.

#### How artifacts differ in ccPAS protocols

The key distinction in ccPAS is the delivery of a second TMS pulse, often at a short inter-stimulus interval, which introduces several challenges:

- **Temporal Overlap:** A critical issue in ccPAS is that artifacts from the first TMS pulse can obscure the early neural response to the second, particularly at short ISIs. Rogasch et al. (2013) showed that even single pulses produce substantial artifacts within the first ∼12 ms, with secondary muscle artifacts—especially from lateral scalp stimulation—lasting up to 40 ms. While these pose challenges for paired-pulse protocols, the authors note that artifact-free EEG signals are generally achievable from ∼10–12 ms post-stimulation when muscle activation is minimized.
- **Amplified Artifacts:** Delivering two pulses increases the complexity of artifact profiles: **Muscle Artifacts:** Each pulse can independently activate scalp and facial muscles, producing distinct bursts of EMG activity. This becomes more problematic when coil positions are close together, as residual muscle activity from the first pulse may overlap with the second. Artifact burden is also location-dependent—lateral frontal sites more often evoke overt muscle twitches than medial frontal regions (Conde et al., 2019). **Ringing and Electrical Artifacts**: The electromagnetic pulse from each coil generates a large, transient EEG artifact ("ringing"). In ccPAS, the presence of two pulses leads to overlapping artifacts, often resulting in a more complex contamination pattern. **Recharging Artifacts:** Some stimulators produce a secondary spike in EEG due to capacitor recharging after each pulse. In paired-pulse protocols, two such recharging artifacts may occur, further complicating isolation of early cortical responses (Hernandez-Pavon et al., 2023).

#### Mitigating ccPAS-Related Artifacts

Artifact reduction in ccPAS protocols requires combining hardware optimization, experimental controls, and advanced post-processing. No single method is sufficient; a multimodal approach is most effective.

1. Hardware and Experimental Setup Strategies

- **Coil Placement:** Precise coil positioning is essential to minimize muscle artifacts in TMS-EEG recordings. Placing the coil away from superficial facial and neck muscles can significantly reduce EMG contamination. Real-time monitoring of TEPs allows on-the-fly adjustments to coil orientation, minimizing craniofacial muscle activity (Casarotto et al., 2022).
- **Auditory Masking:** The coil click evokes auditory potentials that can confound results. Masking noise via sound-attenuating earphones (e.g., ER3C, Etymotic Research) can mitigate this. A sophisticated tool for generating this noise is the TAAC (TMS Adaptable Auditory Control) software (Russo et al., 2022). TAAC can create customized masking noises in real-time for various TMS-EEG protocols, including single pulse, paired pulse, or burst stimulation. The tool uses the stimulator’s specific click sound, tailoring it to an individual’s auditory perception by mixing and manipulating both click and white noise components in the time and frequency domains. This results in noises that are effective at lower, safer volumes compared to existing methods.
- **Electrode and Amplifier Selection:** TMS-compatible electrodes and amplifiers are engineered for rapid recovery from pulse artifacts. These systems minimize ringing artifacts and allows recovery of early evoked components (Ilmoniemi and Kicić, 2010; Hernandez-Pavon et al., 2023).
- **Stimulation Intensity:** Use the lowest stimulation intensity that still produces the intended physiological effect to minimize peripheral (muscle/nerve) activation and reduce artifact amplitude.
2. Software and Data Analysis Strategies

- **Independent Component Analysis (ICA):** ICA is a widely used technique that decomposes EEG signals into independent sources, enabling the removal of components linked to muscle activity, eye movement, and TMS-induced electrical noise. When applied carefully, ICA can substantially improve signal quality by isolating artifact-related components while preserving genuine neural responses (Korhonen et al., 2011; Rogasch et al., 2014; Hernandez-Pavon et al., 2023).
- **Signal Space Projection (SSP):** SSP uses the spatial structure of artifacts to suppress them while preserving genuine neural signals. It has proven effective in both simulations and empirical studies for removing muscle artifacts without distorting early TEPs (Mutanen et al., 2016, 2022; Hernandez-Pavon et al., 2023).
- **Artifact Interpolation:** When artifacts are too large—especially within the first few milliseconds—recovering genuine neural activity is often infeasible. A common approach is to blank or interpolate these intervals using adjacent data. In ccPAS protocols, interpolation may be required around both pulses to mitigate cumulative artifact effects (Ilmoniemi & Kičić, 2010; Mutanen et al., 2016; Hernandez-Pavon et al., 2023).
- **Sophisticated Modeling:** Advanced modeling approaches—such as quantitative physical models of the electrode-gel-skin interface and neural network-based predictors—are increasingly being used to more accurately characterize and remove TMS-induced artifacts, even within the first few milliseconds post-stimulation. These methods make it possible to recover early neural signals that would otherwise be obscured by artifact decay dynamics or electrical discharge (Freche et al., 2018).
- **Template Subtraction:** This method involves subtracting a non-neural artifact template from the recorded EEG. Here are two approaches: **Sham Subtraction:** A robust method using EEG recorded during sham stimulation to serve as a control artifact template. Subtracting this from real TMS data allows the isolation of genuine TMS evoked neural responses (Biabani et al., 2019). This method is highly effective at removing a wide range of non-neural artifacts; however, it requires a separate sham condition, which adds experimental complexity and time. **Trial-Averaged Subtraction:** Alternatively, an artifact template (Tomasevic et al., 2017) can be created by averaging all trials and then subtracting it from each individual trial. This method is effective for removing consistent artifacts and does not require a separate sham protocol; however, it carries the risk of over subtracting genuine, time-locked neural signals, especially if they closely resemble the artifact in timing and amplitude.

## II. Roadmap: Real-time brain-state-coupled ccPAS

Box 1 is a protocol-style roadmap for brain-state-coupled ccPAS implementation, organized into modular steps that can be adopted independently. Alternative strategies are listed for each stage. In the next section, we demonstrate the feasibility with a proof-of-concept experiment implementing selected roadmap strategies and reporting neurophysiological outcomes.

### Box 1. Roadmap for brain-state-coupled ccPAS implementation

#### 1) Identifying the target brain regions

i. Functionally-defined
  - Use individual functional connectivity maps (fMRI, source-reconstructed EEG, MEG) based on regions activated during cognitive tasks
  - Select regions whose enhanced co-activation with TMS correlates with increased performance in selected tasks
  - Determine regions that have the highest brain state marker signal-to-noise ratio (SNR, [e.g., choose locations with the strongest theta SNR if using theta oscillation as a brain state marker])
ii. Anatomically-defined
  - Diffusion MRI (Tractography-Based Targeting):
  - Use offline or real-time white matter tractography to define structurally connected brain regions
  - Structural MRI (Morphology-Based Targeting):
  - Consider scalp-to-cortex distance, as regions deeper from the scalp may receive a weaker induced electric field.
  - Preferentially target gyral crowns rather than sulcal walls, where the electric field is stronger and more perpendicularly aligned with cortical neurons.

#### 2) Defining the target brain state

- Brain states are multidimensional and can be defined behaviorally, cognitively, neurophysiologically (e.g., EEG/fMRI), or biochemically (e.g., neurotransmitter levels), depending on the research context and measurement modality.
- EEG-based neurophysiological markers—particularly neural oscillations—are preferred for brain-state-dependent stimulation due to their high temporal resolution, real-time compatibility, and ability to reflect dynamic cortical excitability and connectivity which enable precise alignment of stimulation to endogenous brain rhythms and enhance protocol individualization.
- Working memory is an example cognitive paradigm suitable for brain-state-coupled stimulation, as it involves dynamic oscillatory activity (e.g., theta/alpha phase), cross-frequency coupling, and evolving network topology measurable via EEG. Other cognitive domains—such as attention, response inhibition, and perceptual decision-making—are also promising, as they engage well-defined neural circuits and oscillatory signatures that can be tracked in real time.
- Key EEG features used across these paradigms include oscillatory phase, within- and cross-frequency coupling, and graph theory network properties (e.g., centrality, segregation, integration)—which dynamically evolve during cognitive engagement.
- Effective brain-state markers must be *(i)* reliably monitored and computed in real time (within milliseconds to ∼1 second), and *(ii)* exhibit sufficient SNR in the target brain region to support accurate, time-locked stimulation.

#### 3) Optimizing dual-site stimulus parameters

##### 3.1. Stimulation order: conditioning (pre) versus test (post) brain regions

i. Functionally-defined
  - Determine the direction of information flow during a cognitive task of interest
  - Compare the effects of stimulating brain regions in one direction versus the other on cognitive task performance (i.e., which direction leads to increased versus decreased performance)
  - Contrast the effects of stimulation order on neurophysiological metrics (e.g., TMS- evoked potential amplitude)
ii. Anatomically-defined
  - Anatomical tracing with anterograde and retrograde tracers on human and non- human primate tissues to determine origin and termination sites
  - Subject-specific white matter tractography to define pre versus post areas and further refine the coordinates of both stimulation sites

##### 3.2. Inter-stimulus interval

i. Functionally-defined
  - Utilizing a combined TMS-EEG approach:
    a. Applying TMS over one cortical area and calculating the time point with maximal stimulus-evoked EEG amplitude in the second region
    b. Empirically testing different ISIs between the first and second pulses and selecting the ISI leading to the greatest change in evoked-EEG amplitude
    c. Compare the effects of varying ISIs on cognitive task performance (i.e., which ISI leads to increased versus decreased performance)
ii. Anatomically-defined
  - Derive the conduction delay between two regions-of-interest from MRI- based diffusion tractography

##### 3.3. Stimulation intensity

- Electric-field modeling using software such as SimNIBS to calculate the induced electric field in the cortical grey matter of each stimulated region
- White matter tractography-based electric field modeling
- Using TMS-EEG, empirically test different combinations of stimulation intensities for the first and second pulses and selecting the intensity generating the strongest change in evoked-EEG amplitude

##### 3.4. Frequency/ inter-trial interval (i.e., time between pulse pairs) and total number of paired pulses

- Utilizing TMS-EEG, determine the dose-response curve testing the effect of multiple number of total pulse pairs on neurophysiological metrics (e.g., TMS- evoked EEG amplitude) basing the initial inter-trial interval on previously used latencies (e.g., 0.2 Hz)
- After selecting the optimal number of paired pulses, the frequency can be further optimized by testing the effect of multiple frequencies on the chosen neurophysiological metric

#### 4) Mitigating ccPAS TMS-related artifacts

- Multimodal approach: effective artifact mitigation in ccPAS protocols requires integrating hardware, experimental, and software-based strategies—no single method is sufficient.

##### 4.1. Preventive strategies during data acquisition

- Coil placement, stimulation intensity, and EEG hardware: Precise coil positioning—particularly avoiding facial and neck muscles—helps minimize EMG contamination. Using the lowest stimulation intensity that still elicits the desired physiological effect reduces peripheral muscle/nerve-related artifacts. Employing TMS-compatible EEG hardware, including low-impedance electrodes and fast-recovery amplifiers, further improves signal quality.
- Real-time TEP monitoring: Visualizing TMS-evoked potentials (TEPs) during acquisition allows on-the-fly adjustments to coil orientation and target site, helping to maximize SNR and proactively minimize artifacts before they contaminate the data.
- Auditory masking: The TMS coil click induces auditory-evoked potentials that can confound EEG signals. Tools such as TAAC (TMS Adaptable Auditory Control) generate individualized masking noise in real time to reduce this effect. For optimal delivery, insert earphones with high passive attenuation (e.g., ER3C insert earphones from Etymotic Research) are recommended, as they effectively block external sound while delivering masking stimuli at safe and adjustable volumes.

##### 4.2. Real-time artifact suppression

- SOUND (Source Estimate Utilizing Noise Discarding): SOUND is a spatial-filtering method capable of real-time artifact suppression with sub-millisecond latency. It suppresses EMG and stimulation-related artifacts while preserving genuine neural signals—making it suitable for closed-loop EEG–TMS applications.

##### 4.3. Post-processing: Signal decomposition techniques

- Independent Component Analysis (ICA): ICA separates EEG signals into statistically independent components, enabling identification and removal of artifact-related sources (e.g., eye blinks, muscle artifacts, pulse artifacts) while preserving neural activity.
- Signal Space Projection (SSP): SSP is a spatial filtering method that suppresses artifacts based on their characteristic spatial topographies.
- Artifact interpolation: When artifact contamination is too severe—particularly in the first few milliseconds post-TMS—data segments are blanked and replaced using surrounding data. In ccPAS, interpolation is often needed around both pulses.

##### 4.4. Post-Processing: Template-based and model-driven correction

- Template Subtraction:
- Sham-based: EEG recorded during sham stimulation is subtracted from real trials to remove artifacts while preserving neural signals.
- Trial-averaged: An artifact template is created by averaging across trials, then subtracted from each trial. Effective for consistent artifacts, but risks over-subtracting genuine, time-locked neural responses.
- Modeling Approaches: Advanced methods (e.g., biophysical modeling of the electrode–gel–skin interface or neural network predictors) are emerging to reconstruct artifact-free EEG at very early latencies, allowing access to previously obscured early TEP components.

#### 5) Other experimental considerations

- Include a sham control condition to isolate the true effects of stimulation on cognitive performance, accounting for both baseline variability and placebo-related influences.
- Since no sham condition fully replicates the multisensory experience of real TMS—particularly the auditory click and scalp sensation—an active cortical control site can help disentangle neural effects from non-specific stimulation artifacts.

## III. Real-time fronto-parietal brain-state-coupled ccPAS

### III.A Experimental design

The experiment was approved by the local ethics committee of the University of Tübingen medical faculty. Seven healthy subjects (3 females, 18-50 years old, mean age: 29 ± 5.5), were tested after written informed consent was provided. Participants recruited had no history of neurological or psychiatric conditions and were not taking any medications affecting the central nervous system. Participants were instructed to adhere to their usual daily routines, sleep at least 6–7 hours prior to each session, and abstain from consuming alcohol or other psychoactive substances up to 2–3 days beforehand. Individuals with habitual alcohol or nicotine use were excluded. This experiment has a three-arm pseudo-randomized crossover design, with ccPAS targeting prefrontal positive theta peak as the experimental condition and random phase ccPAS as the primary control. Each subject was randomly assigned to start with either theta-coupled (POS) or uncoupled (RAND) ccPAS, then they underwent the other condition at least 1 week after the first session. At the reviewer’s request, theta phase-locked prefrontal (PREF) control was added. PREF used the same ccPAS delivery algorithm with a Sham parietal pulse (coil perpendicular to the scalp), effectively stimulating only the dmPFC target.

The experiment began with a 7-minute resting state EEG recording (Figure 4), which was analyzed to extract theta activity from the dmPFC using an individual source-based spatial filter. Source-based spatial filters were constructed from subject-specific MRI scans prior to the experiment. From the individual T1-weighted anatomical sequence, we segmented and meshed tissue boundaries, constructed a three-compartment boundary element (BEM) volume-conductor head model, and computed the corresponding EEG forward solution using a custom pipeline (Stenroos and Sarvas, 2012; Stenroos and Nummenmaa, 2016). For individual source reconstruction, we employed a cortical mid-thickness surface, defined as the geometric mean of the white- and pial-surface boundaries. The source space consisted of ∼16,000 vertices. Each vertex was adjusted to a spherical template to preserve sulcal–gyral topology representation. This procedure optimized vertex-wise correspondence across participants and ensured approximately uniform sampling of the cortical mantle while maintaining individualized modeling. Electrode positions, pinpointed on each subject’s head using the Localite neuronavigation system and projected onto the scalp mesh, were used to define the sensor geometry for the EEG forward solution (lead field matrix), which specifies the predicted voltage topographies across electrodes for each cortical dipole. To estimate theta source activity, the resting-state EEG recording was projected into source space for spectral analysis. Starting from an initial dmPFC MNI seed, the experimenter iteratively sampled neighboring coordinates. At each candidate site, the nearest 20 vertices were selected and their corresponding lead field vectors—together with a regularized sensor covariance—were used to derive local linear constrained minimum variance (LCMV) beamformer spatial filter weights (Van Veen et al., 1997) and project a dmPFC time series. The spectrum was then estimated after aperiodic removal using Irregular Resampling Auto-Spectral Analysis (IRASA) (Wen and Liu, 2016), and theta SNR was calculated (Donoghue et al., 2020). The coordinate yielding the maximal theta SNR was selected to define the final subject-specific dmPFC spatial filter, and dipoles within a 1-cm diameter centered on this coordinate were defined as the dmPFC stimulation target.

After defining the dmPFC ROI with the highest theta SNR, we proceeded with the intervention which consisted of Sham (125 pulse pairs) and Real (250 pulse pairs) ccPAS blocks. The ccPAS protocols were applied while subjects were at rest. During the Real ccPAS block, TMS pulse pairs were applied every ∼3 seconds through the real-time system (bossdevice, sync2brain, Germany) when pre-defined features of the theta oscillation were met. In addition to the real-time phase estimation from ongoing EEG data, other constraints were included to take into account the presence of artifacts and the intrinsic fluctuations characteristic of theta oscillations (Gordon et al., 2021). Thresholds were adjusted incrementally, as needed, during the stimulation period: *1)* eye movement—which blocks TMS triggers for the following 700 ms after an eye blink was identified; *2)* general EEG artifact (including muscle artifact)—to ensure magnetic stimulation is applied in the absence of high-amplitude artifacts; *3)* theta phase stability—to only send a trigger if no phase reset was detected in the last 500 ms; and *4)* theta amplitude—to send a trigger only when a reliable theta oscillation of sufficient amplitude was present. Initial thresholds for all parameters were pre-calibrated immediately before the Sham block. While running the stimulation script, thresholds were adjusted such that pulse pairs reliably triggered at a consistent inter-trial interval (ITI) of 3 seconds (0.33 Hz). This ensured that thresholds were calibrated under realistic conditions, before any stimulation was delivered to the cortex. The script included a hard-coded minimum time constraint of 3 seconds. Thus, even if all gating criteria were met, no pulse pair could be delivered more frequently. These methods were validated by previous work in our lab and have been reported to increase phase detection accuracy (Gordon et al., 2021).

The real-time phase decoding process was identical in both Real and Sham blocks; however, during Sham, the TMS coils were placed perpendicular to the surface of the target regions such that no cortical electromagnetic induction occurs. To control for potentials in the EEG response representing peripherally evoked potentials (PEPs) (Conde et al., 2019), we applied masking noise to suppress auditory evoked potentials from the TMS “click” sounds as well as electrical stimulation to control for somatosensory evoked potentials from the tactile sensation felt during each TMS pulse. This procedure and the subtraction of signal during Sham stimulation from the Real TMS block have been demonstrated to effectively remove PEPs from the direct cortical response to TMS (Gordon et al., 2021).

### III.B Implementation of Roadmap Steps

#### 1) Identifying t arget brain regions

We targeted a two-node FP circuit supporting visual-spatial working memory, defining individualized dmPFC and parietal nodes using convergent functional evidence and anatomical constraints. Based on prior visual-spatial working memory literature, we selected FPN subregions reliably engaged by the task.

The individualized dmPFC stimulation site was then identified via a manual, GUI-guided local LCMV beamforming search for the peak theta SNR within dmPFC. A custom interactive interface leveraged each participant’s MRI-derived head model (BEM surfaces and source mesh), realigned electrode montage (NAS/RTR/LTR), subject-specific lead field, and covariance estimated from 3-s resting-EEG segments. Beginning from an initial dmPFC MNI seed, the operator iteratively sampled nearby coordinates; at each candidate site, the tool selected the nearest N vertices (default N = 20), computed a local LCMV spatial filter with a regularized covariance (λ = 0.001·max eig(Cov)), projected a virtual dmPFC time series, and estimated the oscillatory spectrum using IRASA (aperiodic removed). Theta-band SNR (4–8 Hz) was quantified at each location, and the maximal-SNR coordinate was exported in MNI space for neuronavigation.

The superior parietal lobule (SPL) target was determined visually based on individual T1-weighted MRI. We targeted the SPL–precuneus junction, centering the ROI on the medial lip of BA7 at the dorsal (superior) edge of the IPS as it turns onto the medial wall, posterior to the postcentral sulcus, anterior to the parieto-occipital sulcus, and superior to the subparietal/marginal cingulate sulcus. The placement sat on the medial SPL gyral crown—capturing the border zone between lateral BA7 (7A/7P) and the medial precuneus (7m)—while avoiding the deep medial wall.

For both prefrontal and parietal targets, we also constrained targeting using individual structural MRI in the neuronavigation system to favor gyral crowns and to align the coil such that the induced current ran approximately perpendicular to the local gyral crest (Opitz et al., 2011; Thielscher et al., 2011; Bungert et al., 2017; Laakso et al., 2018; Lynch et al., 2022; Klomjai et al., 2015). TMS-induced electric fields concentrate at gyral crowns and attenuate in sulcal walls; moreover, orienting the coil to drive current orthogonal to the gyrus maximizes the normal field component that most effectively depolarizes pyramidal elements and reduces inter-subject variability and intensity demands relative to suboptimal orientations (Thielscher et al., 2011; Opitz et al., 2011; Laakso et al., 2018; Siebner et al., 2022; Deng et al., 2013). This morphology-informed placement improves focality and consistency compared with stimulating sulcal banks, where greater depth and unfavorable geometry yield weaker, less reliable recruitment (Opitz et al., 2011; Deng et al., 2013; Bungert et al., 2017; Lynch et al., 2022). The Localite neuronavigation system was used to ensure that the location and orientation of the two TMS coils remained constant throughout the stimulation session.

#### 2) Defining the brain state marker

Oscillation phase provides a temporal window for input convergence (Buzsáki and Draguhn, 2004). Further, different phases of an oscillation represent distinct excitability states and applying magnetic or electrical stimulation at specific phases result in differential plasticity effects as demonstrated in humans (Zrenner et al., 2023, 2018; Gordon et al., 2022; Salimpour et al., 2022; Baur et al., 2020) and animal models (McNamara et al., 2022; Escobar Sanabria et al., 2020; Zanos et al., 2018). dmPFC theta was used as the brain state marker in this study due to extensive evidence linking frontal theta activity and FP theta synchrony to working memory and cognitive control processes (Jacob et al., 2018; Heusser et al.,2016; Hsieh and Ranganath, 2014; Phillips et al., 2014; Raghavachari et al., 2001).

Prefrontal theta phase was estimated in real-time from the ongoing EEG signal using the *phastimate* algorithm (Zrenner et al., 2020). Prior to the formal experiment, pilot analyses confirmed that the real-time phase estimation pipeline triggered TMS with sufficient timing precision to reliably target the intended theta phase. As previously described, a subject-specific spatial filter—which incorporated participant anatomy and individualized electrode positioning—was constructed pre-intervention to isolate neural signals originating from the left dmPFC. During the intervention, the personalized source-based filter was applied to ongoing multi-channel EEG to yield a single dmPFC-projected time series. The filtered signal is then the only signal passed into the real-time phase estimation algorithm. The dmPFC-projected time series was downsampled to 250 Hz. Then the most recent 1,024-ms sliding window (256 samples) was bandpass-filtered (5-8 Hz) and trimmed by removing the 35 samples closest to the marker to reduce filter edge artifacts. That data segment was used for forward prediction using a Yule-Walker autoregressive model (Chen et al., 2013), with a total predicted interval of 268 ms (to cover the removed edge and extend 128 ms into the future). A Hilbert transform was subsequently applied to the forecasted signal to obtain an instantaneous phase and amplitude (envelope) estimate, updated every 4 ms. Triggering of both TMS stimulators occurred via an auto-generated TTL pulse when phase and amplitude met preset criteria, the inter-trial interval was 3 s, andno high amplitude artifacts were detected (Figure 3).

**Figure 2.**
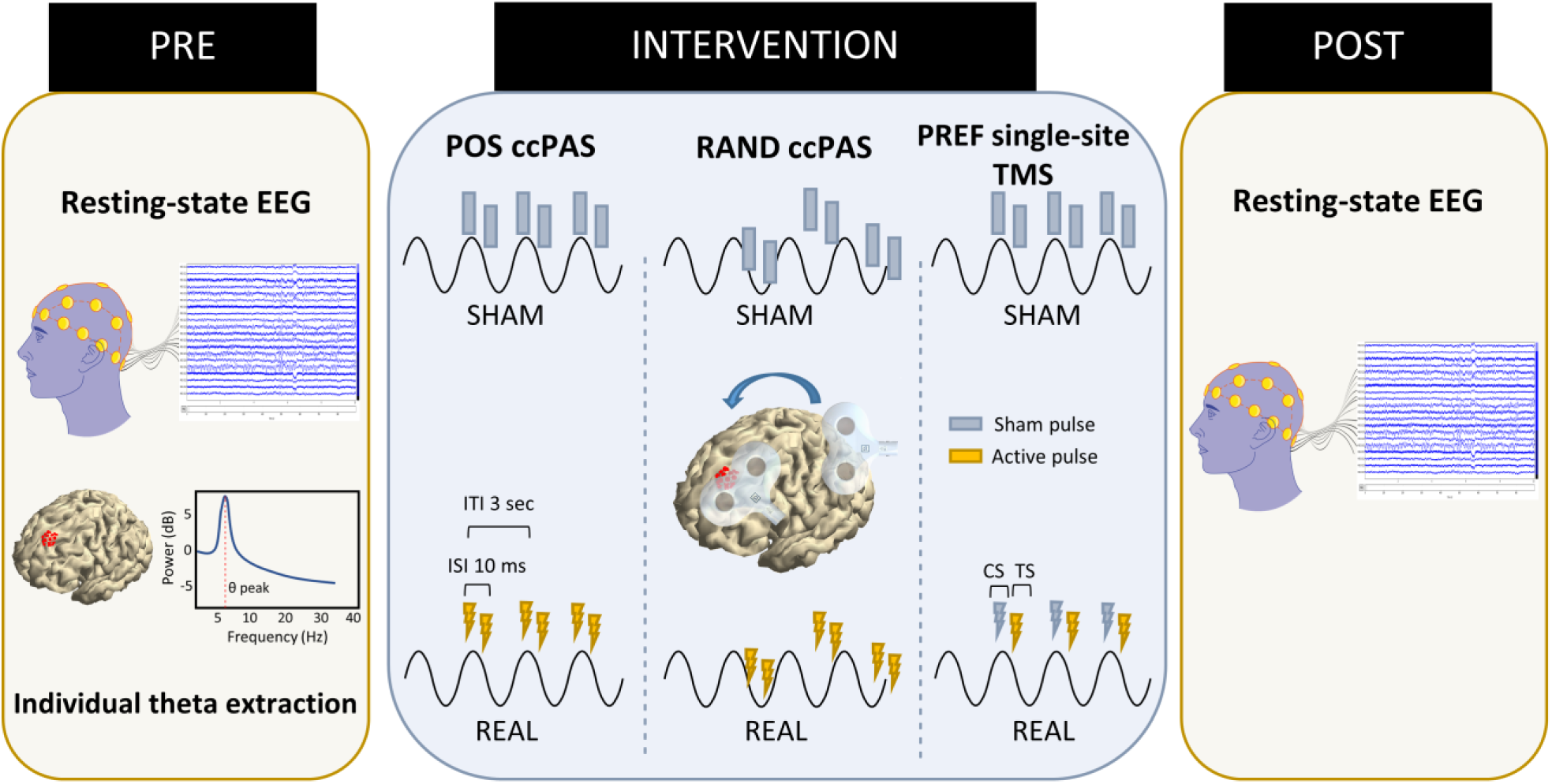
Experimental design. Prior to the main experiment, we extracted prefrontal theta activity from resting-state EEG and performed source-level spectral analysis to identify the medial prefrontal subregion with the highest theta SNR for each individual. This process functionally identified the prefrontal stimulation site, while the parietal target was anatomically defined from individual T1-weighted MRI. Participants were randomized to brain-state-coupled (POS; ccPAS at theta positive peak) or uncoupled (RAND; ccPAS at random theta phase) stimulation, while the alternate protocol administered ≥1 week later. A theta phase-locked prefrontal (PREF) was added as an additional control. Each intervention began with a Sham block (125 pulse pairs), followed by a Real block (250 pulse pairs). Stimulation was delivered from superior parietal lobule to medial prefrontal cortex. Coil placement for prefrontal and parietal targets is illustrated. ccPAS, cortical paired associative stimulation; ISI, inter-stimulus interval; ITI, inter-trial interval; CS, conditioning (first) stimulus; TS, test (second) stimulus

**Figure 3.**
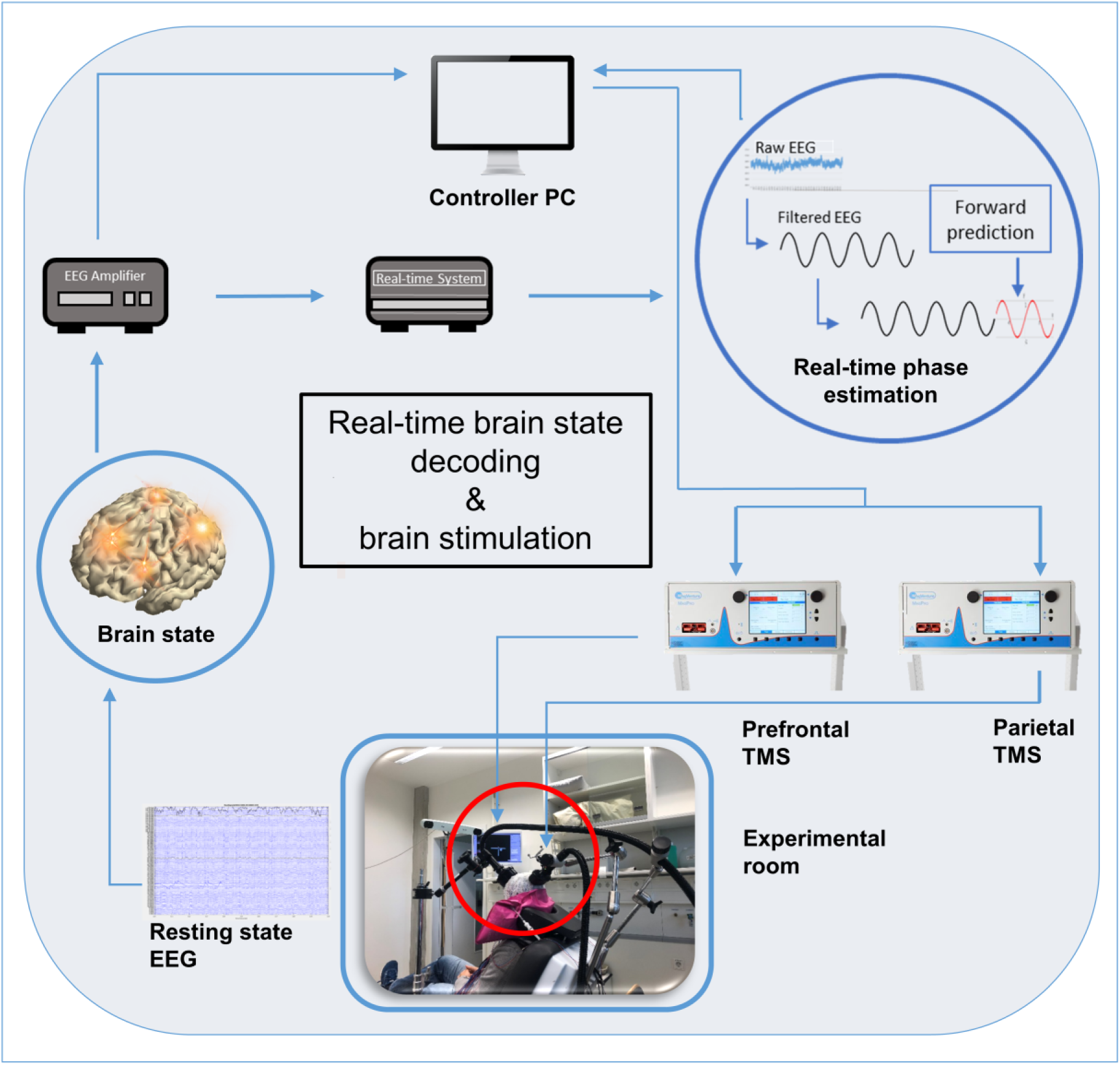
Brain-state-dependent stimulation system. The system integrates real-time EEG recording, processing, and detecting of theta oscillatory characteristics (phase, power, instability) as markers of brain states using a real-time system which then triggers two TMS stimulators when pre-set criteria are met.

#### 3) Optimizing stimulus parameters

##### Stimulation order and inter-stimulus interval

We stimulated SPL → dmPFC with a 10 ms inter-stimulus interval. A 10 ms pairing falls within a commonly used causal STDP window in rodent slices (Anisimova et al., 2022; Jackson, 2020), while human cortical slices show LTD at 5–10 ms that can switch to LTP under dopaminergic tone (Louth et al., 2021; Verhoog et al., 2013). We chose 10 ms to match prior FP ccPAS in humans (Momi et al., 2020; Nord et al., 2019; Santarnecchi et al., 2018) and enable cross-study comparability. Coil orientation at both sites was adjusted to drive current approximately orthogonal to the local gyral crest (see targeting section).

##### Intensity

ccPAS TMS was delivered using two MagPro stimulators (MagVenture A/S, Denmark), each connected to a 46 mm figure-of-eight coil (Cool-B35 HO) delivering biphasic pulses. Each pulse was applied at 130 or 140% of resting motor threshold (RMT) such that the maximum stimulator output (MSO) is > 85%. RMT was defined as the lowest stimulus intensity (%MSO) that evoked an MEP of ≥50 μV in at least 5 out of 10 consecutive trials in the relaxed first dorsal interosseous (FDI) or abductor pollicis brevis (APB) muscle of the right hand (Rossini et al., 2015). TMS coils used in the current experiment were smaller than previous ccPAS studies which used 70 mm figure-of-eight coils at 120% RMT (Casula et al., 2016; Santarnecchi et al., 2018; Nord et al., 2019; Momi et al., 2020). Other studies also used a multi-channel deep TMS system with two H-D1 coils (Zibman et a., 2019; Aviram-Friedman et al., 2025) which penetrate 1.5-2x deeper than conventional coils. High intensities were applied in the current study to ensure activation of prefrontal and parietal targets which are located deeper than the M1-hand area used to define the RMT.

##### Frequency and number of pulse pairs

The frequency between pulse pairs was ∼0.33 Hz (ITI ≈3 s). While previous ccPAS TMS protocols have used paired stimuli at lower repetition frequencies (0.1 to 0.25 Hz), STDP has been reliably induced across a broader range of paired repetition frequencies—from 0.1 Hz up to 10 Hz—using various stimulation modalities, including TMS, electrical, and optogenetic stimulation. These protocols have been validated in brain slices, *in vivo* preparations, and human studies (Turrini et al., 2024, Borgomaneri et al., 2023, Anisimova et al., 2022, Inglebert et al., 2020, Seeman et al., 2017, Sgritta et al., 2017, Campanac & Debanne, 2008). In addition to repetition frequency, the number of pulse pairs is also an important design consideration. Previous ccPAS studies employed ∼100-180 paired stimuli (e.g., Momi et al., 2020; Nord et al., 2019). Cui et al. (2018) found that increasing STDP pairings from 100 to 250-300 pairs increased the robustness of NMDAR-dependent LTP against jitter. To account for possible temporal variability during phase-locked stimulation, we applied 250 pulse pairs to increase the robustness of LTP induction.

#### 4) Mitigating artifacts

We mitigated TMS-related artifacts using a combined acquisition-control and post-processing approach. First, to suppress auditory-evoked potentials from the coil click, participants wore insert earphones with high passive attenuation (e.g., ER3C), and masking noise was delivered and adjusted to perceptually eliminate the click (generated with TAAC; Russo et al., 2022). Second, we implemented an optimized sham block (Gordon et al., 2021) that preserved identical timing and phase-triggering but rotated the coil perpendicular to the scalp to prevent cortical induction and added time-locked cutaneous electrical stimulation to approximate the tactile sensation of real TMS; sham recordings were later used as a non-neural template.

In post-processing, brief pulse-contaminated segments surrounding each conditioning and test pulse were blanked and interpolated to remove saturating decay/charging transients while preserving later TEP components (Mutanen et al., 2016; Hernandez-Pavon et al., 2023). Independent Component Analysis (two-pass ICA) was then applied to remove components reflecting residual craniofacial EMG, eye blinks/saccades, and auditory responses (Rogasch et al., 2017). For details on EEG preprocessing, please see the section on outcome measures. In addition to cortical responses to TMS, TEPs are contaminated by peripherally evoked potentials (PEPs), which arise from multisensory inputs associated with TMS delivery and produce EEG deflections time-locked to the TMS pulse but not directly attributable to electromagnetic cortical activation (Farzan and Bortoletto, 2022; Hernandez-Pavon et al., 2022). To attenuate PEPs resulting from auditory and somatosensory inputs, sham-template subtraction (Real − Sham) was performed on a within-subject basis (Biabani et al., 2019; Gordon et al., 2021).

Although the resulting difference waveform is commonly interpreted as reflecting EEG responses driven primarily by direct cortical activation, this interpretation warrants caution. Sham subtraction assumes linear superposition between neural and non-neural (peripheral) components—an assumption that may not always hold. Auditory, somatosensory, and muscle-related inputs may interact nonlinearly with ongoing cortical dynamics and distort genuine neural responses (Conde et al., 2019). Moreover, artifact suppression techniques during preprocessing may differentially affect Real and Sham conditions, particularly at early latencies dominated by large-amplitude artifacts (Rocchi et al., 2021; Biabani et al., 2019). Despite these limitations, sham subtraction remains a widely used and practical control approach when closely matched for coil orientation and multisensory stimulation (Conde et al., 2019; Rocchi et al., 2021). Accordingly, sham subtraction was applied conservatively in the present study and complemented with additional artifact-mitigation procedures to maximize signal fidelity.

#### 5) Defining outcome measures

To evaluate the effects of brain-state-coupled and uncoupled ccPAS on neural dynamics, we examined three complementary outcome measures: *(1)* TMS-evoked potentials (TEPs) during ccPAS to probe immediate cortical responses (ccPAS-TEP), *(2)* resting-state EEG functional connectivity before and after stimulation to assess network-level changes, and *(3)* post hoc tractography to provide anatomical context for potential structural pathways mediating stimulation effects.

##### TMS-evoked potentials (TEPs)

TEPs consist of a typical sequence of deflections (peaks and troughs) in EEG at specific time points after the TMS pulse (Rogasch et al., 2020; Ilmoniemi & Kičić, 2010). While investigations continue to examine the physiologic properties underlying TEPs, it is widely used as a tool for assessing cortical excitability in the stimulated region as well as the propagation of activation to distant sites in healthy and clinical populations (Belardinelli et al., 2021; Rogasch et al., 2020; Tremblay et al., 2019). For ccPAS-TEP over dmPFC, EEG was epoched −1000 to +1000 ms around the second pulse and averaged channel-wise.

EEG data were preprocessed using a custom pipeline adapted from the methodological framework of the TESA toolbox for TMS-EEG (Rogasch et al., 2017), implemented with FieldTrip (Oostenveld et al., 2011), EEGLAB (Delorme & Makeig, 2004), and custom functions. The pipeline followed core TESA stages: trial segmentation, baseline correction, artifact interpolation (e.g., TMS decay artifact removal), robust detrending, visual inspection and rejection of noisy trials/channels, and two rounds of ICA to remove early and late artifacts (including muscle activity, blinks, auditory-evoked potentials, and line noise).

The combined steps of robust detrending, trial-wise visual rejection, and targeted ICA-based removal were sufficient for addressing muscle artifacts. The first round of ICA was optimized to isolate short-latency, high-amplitude deflections characteristic of TMS-evoked EMG activity. Notably, muscle artifacts were minimal in this study, as stimulation targeted the dmPFC—a para-median site that induces little to no overt muscle twitches, unlike more lateral frontal targets (Conde et al., 2019). Decay artifacts were mitigated using polynomial fitting and exponential modeling tailored to post-pulse decay dynamics. Finally, missing or rejected channels were interpolated via spatial spline methods, and all data were re-referenced to the common average.

#### Resting-State Functional Connectivity

##### wPLI computation across frequency bands

We estimated functional connectivity from 7-min eyes-open resting recording collected pre- and post- intervention using the debiased weighted phase-lag index (wPLI), which reduces volume-conduction bias by emphasizing non-zero-lag phase relationships (Vinck et al., 2011). Band-limited connectivity was computed in theta, alpha, and low-gamma ranges, yielding adjacency matrices for each band. Fronto-parietal connectivity was quantified on frontal and parietal electrodes surrounding stimulation sites. The primary confirmatory ROI included: ‘Fp1’, ‘Fp2’, ‘Fpz’, ‘AFz’, ‘AF3’, ‘AF4’, ‘CPz’, ‘CP1’, ‘CP2’, ‘Pz’, ‘P1’, ‘P2’. A larger ROI graph was also used for supplemental robustness analyses (Table S1; Figure S4).

##### Pre- vs. post-intervention comparisons

For each band, we contrasted pre vs. post connectivity (within-subject) to test whether ccPAS induced durable network-level changes, summarizing effects with region-of-interest means while controlling multiple comparisons via nonparametric permutation. Network diagrams were visualized using BrainNet Viewer (Xia et al., 2013).

###### Post Hoc Tractography of Stimulated Sites

To contextualize possible potential anatomical pathways mediating the observed neurophysiological effects of ccPAS, we performed post-hoc diffusion MRI tractography in a representative participant using DSI studio (Yeh et al., 2013). The individualized dmPFC and SPL targets were used as seeds to visualize tract projections. The diffusion-MRI processing and tractography pipeline is described in Text S1.

## IV. Results

### Brain-state-dependent ccPAS elicits distinct spatio-temporal dynamics of evoked activity

To characterize brain-wide neural response to brain-state-coupled and -uncoupled ccPAS, and to isolate the contribution of phase-locking independent of paired associative stimulation, we examined TMS-evoked EEG responses across conditions. Figure 4 illustrates the sham template subtraction, whereby (A) and (B) show the group-averaged time course and scalp topographies for brain-state-coupled and uncoupled ccPAS, while Figure 5C shows the results for phase-locked prefrontal stimulation. Statistical analyses (Text S2) were done on the cleaned (Real-Sham) signal within time windows of interest (TOI) centered around canonical prefrontal TEP components (P30: 20-35 ms, N45: 35- 50 ms, P70: 50-85 ms (Krile et al., 2023; Ross et al., 2023; Chung et al., 2017). Only signal <100 ms post-TMS was analyzed to minimize contamination from peripheral-evoked potentials (Ross et al., 2022; Biabani et al., 2019). Figure S1 shows the waveforms up to 350 ms post-TMS pulse, while Figure S2 depict scalp maps of the difference between POS ccPAS and the control conditions without sham template subtraction. All statistical tests were whole-head and data-driven, without a priori spatial constraints.

**Figure 4.**
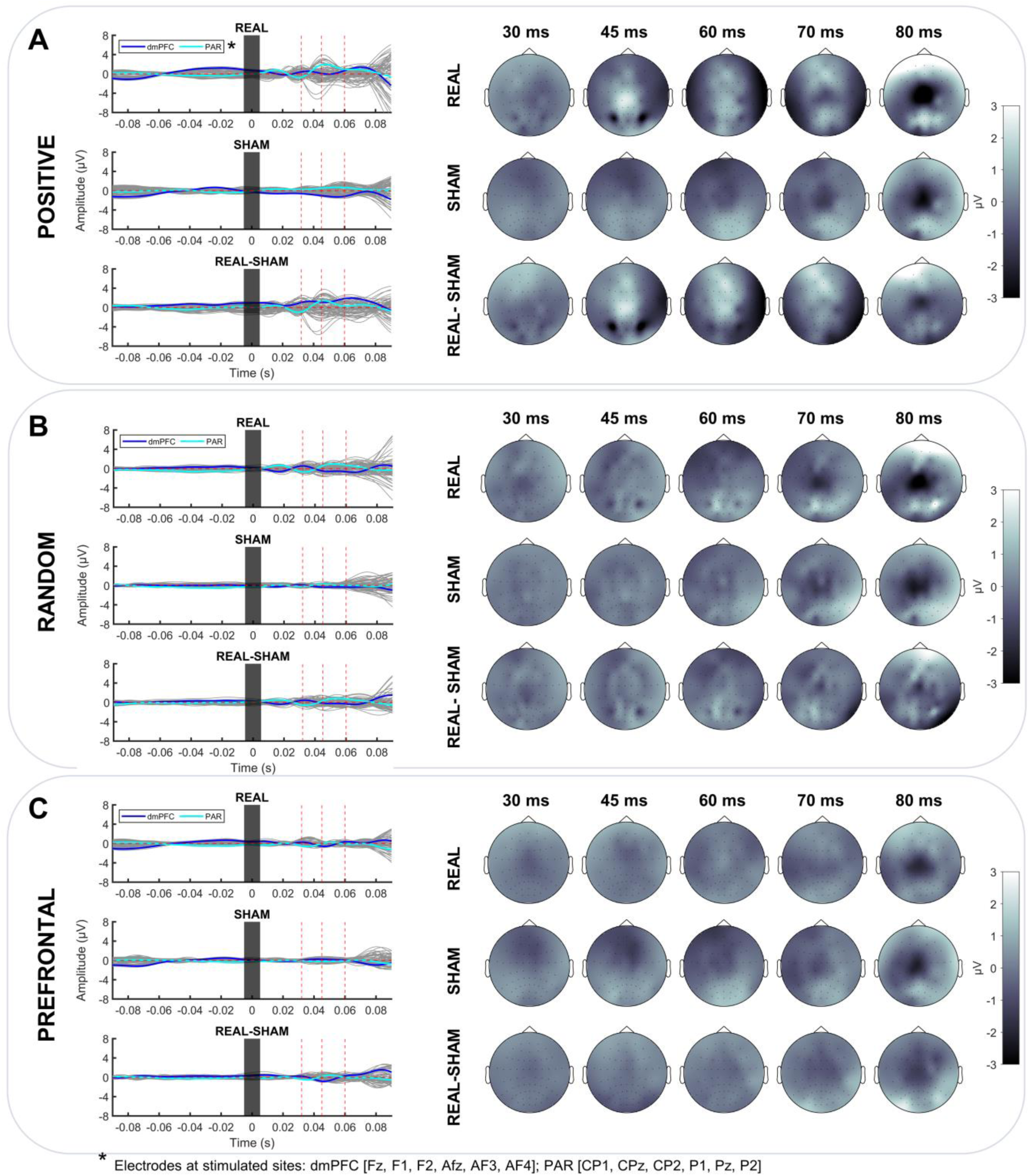
Brain-state-coupled ccPAS of dmPFC and PAR targets induced distinct spatio-temporal changes in brain activity. (A - C) Illustrates sham template subtraction: the process of removing multisensory inputs from direct cortical responses to TMS (Real-Sham). Each section shows the time course (left) and scalp topographies (right) during brain-state-coupled ccPAS [A, Positive], brain-state-uncoupled ccPAS [B, Random], and brain-state-coupled prefrontal only stimulation [C]. This figure is descriptive; corresponding statistical evaluation of these waveforms and topographies are reported in Figure 6. dmPFC, dorsomedial prefrontal cortex; PAR, parietal

**Figure 5.**
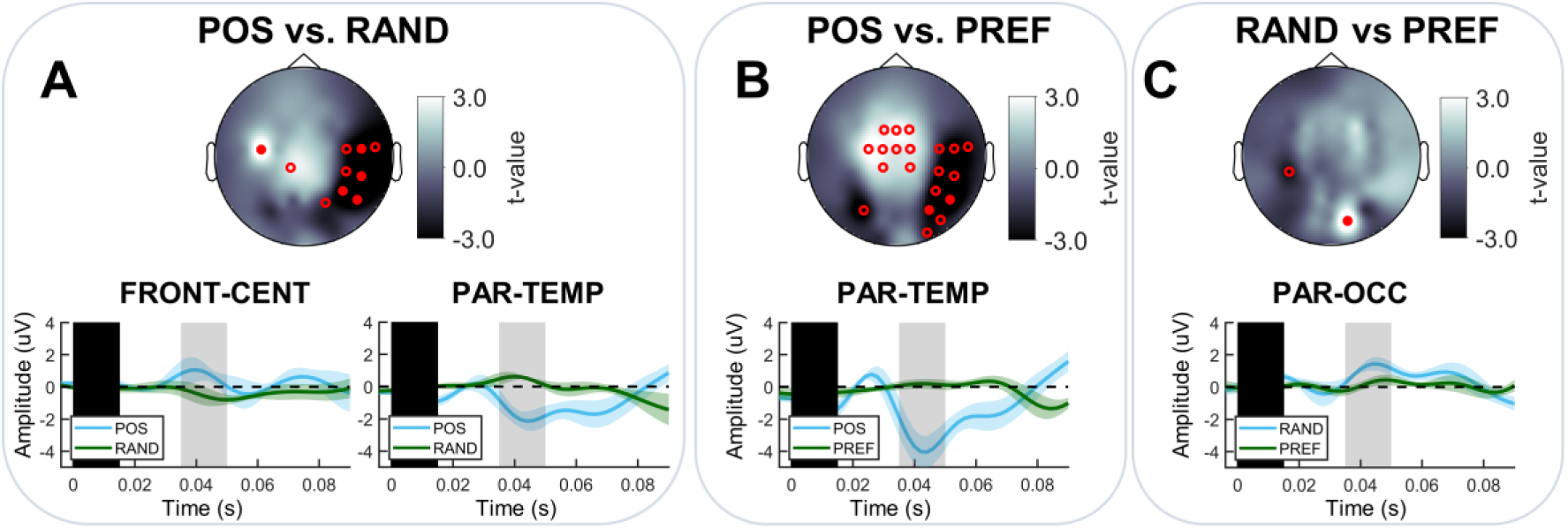
Brain-state-coupled ccPAS produced N45 polarity reversal at fronto-central sensors with right parieto-temporal negativity. (A-C) Confirmatory post-hoc pairwise contrasts within the N45 (35-50 ms) window, defined by the omnibus three-group test. Top: Sensor-level scalp maps show raw GLM t values (fixed scale ±3) across contrasts (POS–RAND, POS–PREF, RAND–PREF). Filled red markers indicate electrodes surviving Holm-FWER correction across contrasts × channels × TOIs; open markers indicate uncorrected p < 0.05. Bottom: ROI-averaged evoked potentials (mean ± SEM) are plotted over contrast-specific significant sensors from scalp maps; condition waveforms are color-coded in each panel. Gray shading denotes the tested interval and the black bar indicates the interpolated TMS-artifact segment. These ROI-averaged waveforms are descriptive illustrations of the sensory group identified by the post hoc tests. POS, phase-locked ccPAS; RAND, phase-uncoupled ccPAS; PREF, phase-locked prefrontal-only stimulation. ROIs: FRONT-CENT, fronto-central; PAR-TEMP, parieto-temporal; PAR-OCC, parieto-occipital

To identify time windows exhibiting condition-dependent modulation, a three group omnibus analysis (POS, RAND, PREF) using a partial-overlap GLM F-test was conducted, with Holm family-wise error correction within each TOI across EEG electrodes. Omnibus and post-hoc statistical inferences were evaluated using Monte Carlo permutation testing implemented in FieldTrip with an exchangeability-preserving resampling method. The omnibus test revealed significant effects exclusively in the N45 latency at right temporal and temporo-parietal electrodes (T8, TP8: F(2,12) = 11.23–14.51; p = 0.0256–0.0441; η²ₚ = 0.65–0.71), indicating very large effects relative to conventional η²ₚ benchmarks (Cohen, 1988; Richardson, 2011; Lakens, 2013). Two additional electrodes showed trend-level effects after Holm correction: O1 in the P30 window (F(2,12) = 5.77, p = 0.070, η²ₚ = 0.49 [very large effect]) and FC5 in the N45 latency (F(2,12) = 7.32, p = 0.074, η²ₚ = 0.55 [very large effect]).

On the basis of this temporally and spatially constrained omnibus effect, confirmatory post-hoc pairwise comparisons were conducted within the N45 interval using partial-overlap GLM t-tests. Whereas within TOI correction across channels was used in the omnibus for temporal localization, confirmatory post-hoc inference was based on a global Holm family-wise error correction across contrasts x channels x TOIs. Effect sizes are reported as r-equivalents derived from GLM t-statistic, with r ≈ 0.1, 0.3, ≥0.5, and ≥0.7 interpreted as small, medium, large, and very large effects, respectively; Cohen’s d thresholds are not directly applicable to r-equivalent or η²ₚ metrics (Cohen, 1988; Rosenthal & Rosnow, 2009; Field, 2013).

### Positive ccPAS produced fronto-central polarity reversal of canonical N45 component with right parieto-temporal negativity

Figure 5 time course and topographical maps illustrate the results of the confirmatory post-hoc tests at N45. In the POS vs. RAND contrast, significant effects were observed at a left fronto-central electrode (FC5: t(6) = 4.32, p = 0.038, r = 0.87 [very large effect]; POS > RAND) and at spatially localized right parieto-temporal sites (CP6, TP8, T8: t(6) = −4.71 to −4.02, uniform p = 0.038, r = 0.85–0.89 [very large effect]; RAND > POS). Comparing POS vs. PREF, significant effects were confined to right parieto-temporal electrodes (TP8, P6: t(9) = −5.16 to −3.20, uniform p = 0.038, r = 0.73–0.86 [very large effect]; PREF > POS). Finally, when testing both control conditions (RAND vs. PREF), a single significant effect was detected at a right parieto-occipital electrode (PO4: t(9) = 5.03, p = 0.038, r = 0.86 [very large effect]; RAND > PREF).

### Exploratory post-hoc tests at P30 window

Given the omnibus trend-level effect in the P30 window (F(2,12) = 5.77, p = 0.070, η²ₚ = 0.49) which signifies a very large η²ₚ effect size (Cohen, 1988; Rosenthal & Rosnow, 2009; Field, 2013), we conducted exploratory post-hoc partial-overlap GLM t-tests to characterize potential condition differences at this latency (Figure 6). Effects surviving Holm familywise error correction was observed only in the POS vs. RAND contrast. Specifically, significant effects were observed at prefrontal electrodes (Fp2, AF3, AF4: t(6) = 2.84–5.28, uniform p = 0.038, r = 0.76–0.91 [very large effect]; POS > RAND) and at a left occipital electrode (O1: t(6) = −2.64, p = 0.038, r = 0.73 [very large effect]; RAND > POS). No Holm-corrected effects were observed for POS vs PREF or RAND vs PREF.

**Figure 6.**
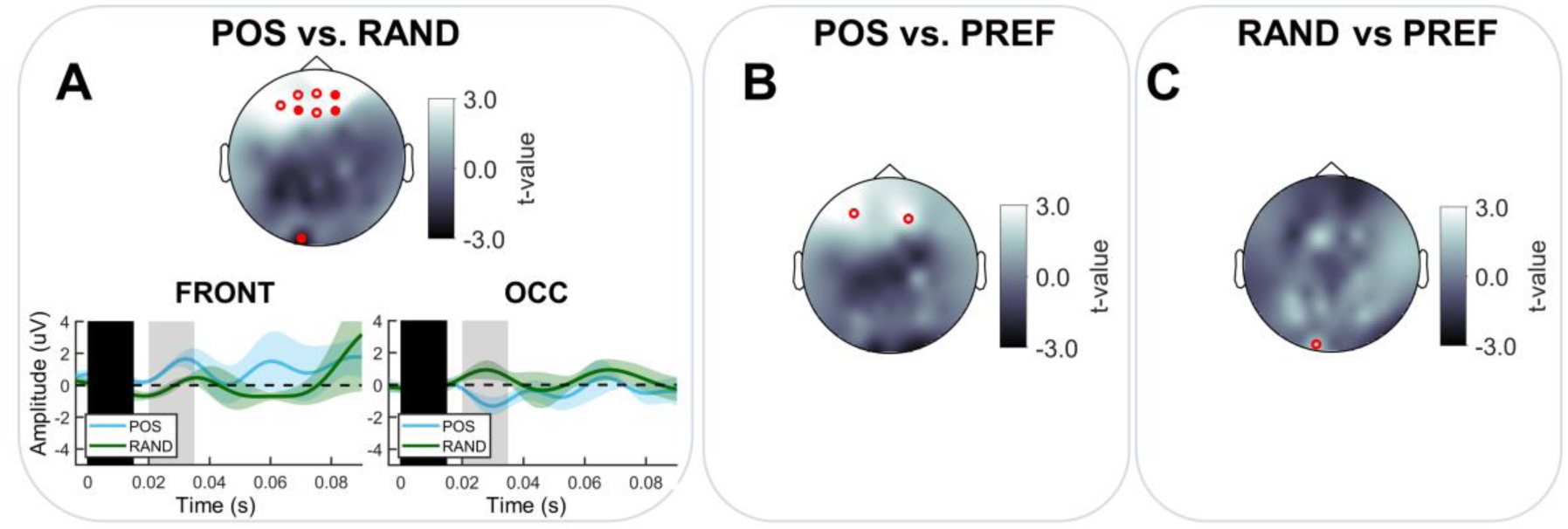
Exploratory post hoc testing at P30 reveal fronto-occipital dissociation for POS vs. RAND. Top: Sensor-level scalp maps show post hoc pairwise contrasts (POS–RAND, POS–PREF, RAND–PREF) in the P30 (20-35) window, motivated by a trend-level omnibus three-group test with a large η²ₚ effect size (F(2,12) = 5.77, p = 0.070, η²_p = 0.49). Maps display raw GLM t values (fixed scale ±3). Filled markers indicate electrodes surviving Holm-FWER correction across contrasts × channels × TOIs; open markers indicate uncorrected p < 0.05. Bottom: ROI-averaged time courses (mean ± SEM) are shown for contrast-specific significant sensors from scalp maps. Holm-FWER–surviving effects were observed only for POS vs RAND. No Holm-corrected effects were observed for POS vs PREF or RAND vs PREF. Condition waveforms are color-coded in each panel. Gray shading denotes the tested interval and the black bar indicates the interpolated TMS-artifact segment. These ROI-averaged waveforms are descriptive illustrations of the sensory group identified by the post hoc tests. POS, phase-locked ccPAS; RAND, phase-uncoupled ccPAS; PREF, phase-locked prefrontal-only stimulation. ROIs: FRONT, frontal; OCC, occipital

To evaluate whether the posterior effect observed at O1 in the P30 window reflected an isolated noisy electrode, spatial and variance diagnostics were done which revealed O1 variability was within the posterior ROI range across subjects (SD = 1.31; MAD = 0.48)—with dispersion values well within the range of a bilaterally defined posterior ROI (median SD = 0.95, range = 0.54–1.41). Critically, the direction of the condition difference at O1 was fully concordant across all posterior ROI electrodes (11/11), indicating a spatially coherent posterior field rather than an isolated sensor artifact. To further assess the robustness of this posterior effect and rule out sensitivity to individual subjects, leave-one-subject-out (LOSO) analyses were performed. The O1 effect remained significant in all folds after Holm family-wise error correction (p ≤ 0.05 in 7/7 folds), with identical p-values across folds (p = 0.013), demonstrating that the effect was not driven by any single subject and was highly stable to sample perturbation.

Altogether, early TMS-evoked activity revealed distinct spatio-temporal response patterns for brain-state-coupled vs. –uncoupled ccPAS and single site control protocols. Confirmatory post-hoc tests showed robust condition-specific modulation in the N45 window over fronto-central and parieto-temporal electrodes. An earlier P30 modulation was also evident in prefrontal and left occipital electrodes only for the POS vs RAND contrast after exploratory analyses. Because TEPs capture instantaneous cortical responses to TMS, we next examined whether these evoked effects translate into sustained changes in FP network connectivity assessed from post-intervention resting state recordings.

### Brain-state-dependent ccPAS reconfigures fronto-parietal connectivity

To demonstrate that the observed changes in EEG activity are not merely transient phenomena associated with phase-locked and/or dual-site stimulation, but rather indicative of genuine network-level modulation, FP connectivity analysis of the immediate post-intervention period was performed. To isolate the effect of phase-locked paired associative stimulation, a POS vs. mean control contrast was implemented as the primary contrast by averaging across control conditions that separately remove phase-locking (RAND) or paired associative stimulation (PREF). Secondary contrasts comparing POS vs. RAND and POS vs. PREF were also evaluated to probe the specificity of the main effect against each control ingredient separately, thereby isolating comparisons to the paired, non-phase-locked and the phase-locked single-site controls. Baseline comparability tests were performed to verify that baseline FP debiased weighted phase lag index (wPLI) levels and variance structure of Δ (post–pre) were comparable across conditions—ensuring that any post-intervention connectivity differences reflect protocol-specific modulation rather than pre-existing group differences or unequal measurement noise. We then quantified whether POS produced larger ΔwPLI changes than the control conditions using a hypothesis-driven, one-sided linear contrast. Edgewise statistic maps were estimated with studentized test statistics for partially overlapping samples (Derrick et al., 2017) and submitted to two permutation-based network inference methods: a family-constrained network statistic (cNBS) adapted from Noble and Scheinost, (2020) and threshold-free network-based statistics (TFNBS; Baggio et al., 2018). These methods capture complementary forms of network reconfiguration: distributed FP changes and edgewise restructuring, respectively. See Text S3 for functional connectivity quantification and statistics.

We hypothesized that the phase-specific FP stimulation in POS would yield a network state different from both controls. Specifically, we predicted that phase-specific ccPAS would induce increases in FP connectivity in theta and alpha, and decreases in gamma. Because ccPAS was phase-locked to endogenous mPFC theta positive peak, we hypothesized that this phase-specific timing facilitates theta synchrony between the stimulated FP areas. Similarly, phase-specific stimulation is known to enhance alpha synchrony when delivered at functionally relevant oscillatory phases (Romei et al., 2016; Thut et al., 2011). Further, stronger alpha coordination is commonly interpreted in top-down inhibitory/gating control (Sadaghiani & Kleinschmidt, 2016; Fries, 2015). Consistent with frameworks in which feedback processing are preferentially carried via alpha rhythms and feedforward processing by gamma (Michalareas et al., 2016; Fries, 2015; van Kerkoerle et al. 2014), increased alpha coordination may coincide with reduced gamma expression, yielding an inverse alpha-gamma relationship (Bonnefond & Jensen, 2025; Jensen, 2024; van Kerkoerle et al., 2017; Jensen & Mazaheri, 2010).

### Baseline FP connectivity exhibits no between-condition bias

Mean PRE FP wPLI in theta, alpha, and gamma bands did not differ across POS, RAND, and PREF conditions (Figure 7A). PRE edgewise t-value distributions for all three contrasts were centered near zero, with only modest band-specific shifts (Figure S3A), indicating no large systematic baseline bias between groups. To assess whether within-subject variability of Δ connectivity differed across conditions, we computed subject-level FP mean Δ and compared variance using Brown-Forsythe tests. Δ variability was comparable across conditions in all bands (Figure S3B). Finally, edgewise Δ-variance structure showed that alpha edges generally had larger Δ variability than gamma edges, but no evidence of extreme or irregular heterogeneity that would threaten exchangeability (Figure S3C). Together, these diagnostics support the exchangeability assumptions required for permutation-based cNBS and TFNBS.

**Figure 7.**
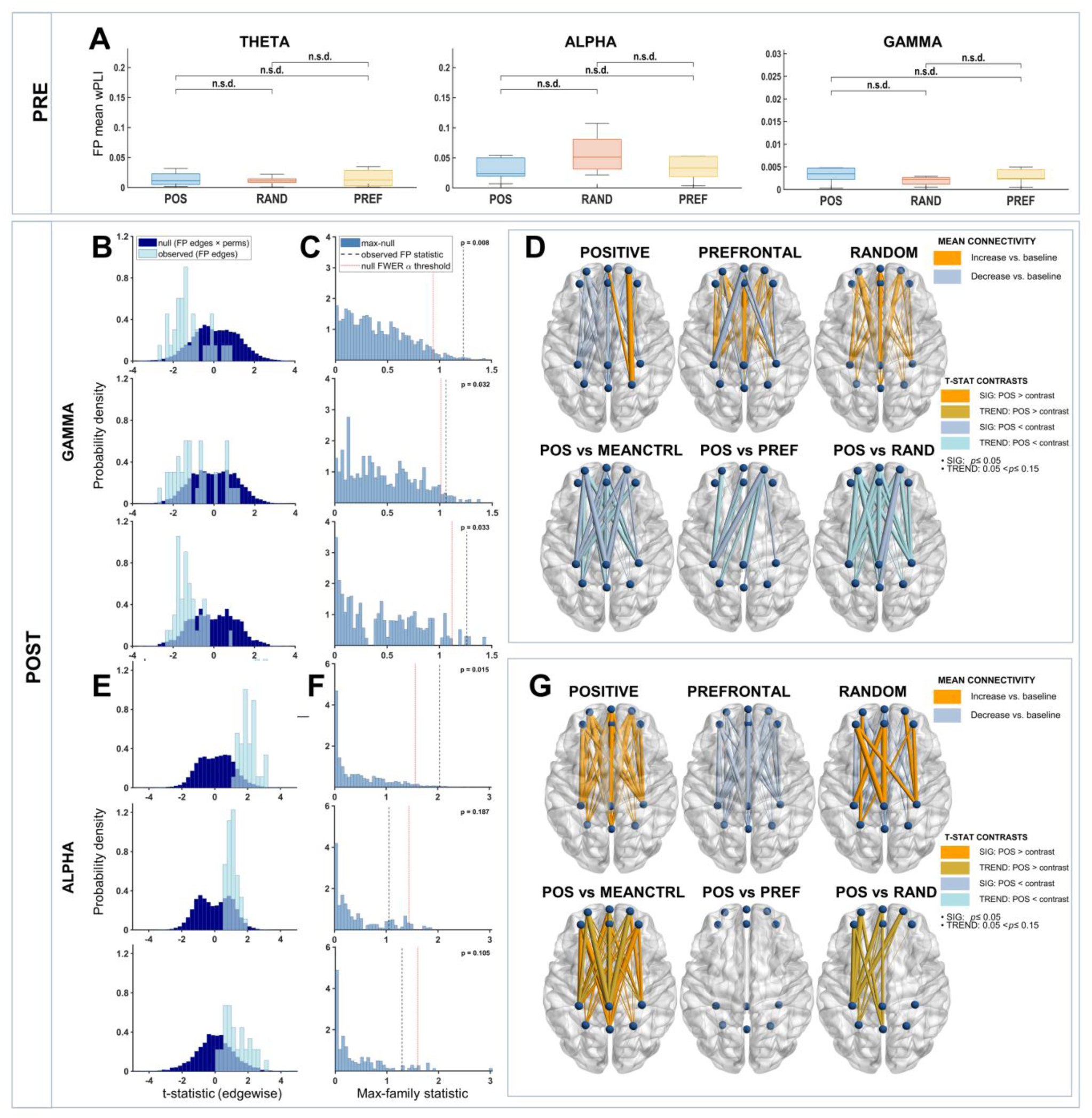
Brain-state-coupled ccPAS drives frequency-specific fronto-parietal reconfiguration: ↓ gamma, ↑ alpha connectivity. (A) Baseline (PRE) fronto–parietal (FP) connectivity quantified with debiased weighted phase lag index (dwPLI) did not differ across Positive (POS, phase-coupled ccPAS), Random (RAND, phase-uncoupled ccPAS), and Prefrontal (PREF, phase-coupled single-site stimulation) conditions in theta (4–8 Hz), alpha (9–12 Hz), and gamma (60–90 Hz) bands. <boxcap>Boxplots show subject-level FP mean connectivity (center line, median; box, interquartile range; whiskers, most extreme non-outlier values). (B–C) Gamma FP network-based inference indicates reduced FP family connectivity after POS. (B) cNBS edgewise t-statistic distributions for FP edges show the null distribution pooled across permutations x FP edges (navy) and the observed FP edge distribution (light blue) [top: POS vs. mean control (average of PREF and RAND); middle: POS vs. PREF; bottom: POS vs. RAND] (C) cNBS max-family null distributions for the FP edge family show family-level effects for POS vs. mean control (p = 0.008), POS vs. PREF (p = 0.032), and POS vs. RAND (p = 0.033) Vertical dashed lines indicate observed family statistics; red dotted line denotes the null FWER α threshold, (D) Gamma FP connectivity diagrams. Top: mean ΔwPLI (POST − PRE) for POS, PREF, and RAND with the top-30 edges illustrated. Bottom: TFNBS edgewise connectivity diagrams show reduced gamma connectivity after POS ccPAS. POS vs. mean control has 14 significant (p = 0.013–0.038) and 8 trend-level (0.052–0.143) edges. POS vs. PREF show 5 significant (p = 0.038–0.048) and 6 trend-level edges (p = 0.053–0.092). POS vs. RAND reveal 3 significant (p = 0.033–0.048) and 20 trend-level edges (p = 0.054–0.149). Darker shades of warm colors represent significant increases (p_FWER ≤ 0.05), while cool colors represent significant decreases (p_FWER ≤ 0.05). Lighter shades denote trend-level effects (0.05 < p_FWER < 0.15). Nodes correspond to frontal and parietal electrodes surrounding stimulation sites. (E–F) Alpha FP network-based inference show increased FP family connectivity after POS. (E) cNBS edgewise t-statistic distributions shown with the same conventions as (B). (F) cNBS max-family null distributions for the FP edge family show family-level increase for POS vs. mean control (p = 0.015), no effect for POS vs. PREF (p = 0.187), and a trend-level effect for POS vs. RAND (p = 0.105) (G) Alpha FP connectivity diagrams. Top and bottom use the same conventions as (D). TFNBS edgewise connectivity diagrams show increased gamma connectivity after POS ccPAS. POS vs. mean control has 26 significant (p = 0.016–0.044) and 10 trend-level (p = 0.053–0.138) edges. POS vs. PREF indicate no edges below threshold, while POS vs. RAND reveal 14 trend-level edges (p = 0.113–0.13). cNBS provides FWER-controlled inference at the FP-family level, whereas TFNBS provides FWER-controlled edge-wise inference. MEANCNTRL, mean control; n.s.d., no significant difference, cNBS, constrained network based statistic; TFNBS, threshold-free network based statistic

### Positive ccPAS decreased gamma FP connectivity

Figure 7 panels B-D show convergent evidence for reduced FP gamma connectivity after POS ccPAS. cNBS (negative tail) indicated robust gamma decreases relative to both control interventions: POS vs mean control (FWER-corrected p = 0.008, d ≈ –1.35); POS vs PREF (p = 0.032, d ≈ –1.20); and POS vs RAND (p = 0.033, d ≈ –0.81). TFNBS further revealed FP with robustly reduced connectivity after Positive ccPAS. POS vs mean control had 14 significant edges (p = 0.013–0.038) exhibiting large decreases in gamma connectivity (median d ≈ –0.96, range –1.51 to –0.57) and 8 trend-level edges (p = 0.052–0.143) with medium-to-large decreases (median d ≈ –0.71, range –0.99 to –0.48). POS vs PREF showed a partially overlapping FP subnetwork with 5 significant edges (p = 0.038–0.048), again showing large effect size reductions (median d ≈ –1.18, range –1.46 to –0.97), plus 6 trend-level edges (p = 0.053–0.092) with medium-to-large decreases (median d ≈ –0.91, range –1.11 to –0.47). POS vs RAND revealed 3 significant edges (p = 0.033–0.048), with medium-to-large decreases in gamma connectivity (median d ≈ –0.90, range –0.91 to –0.77), plus 20 additional trend-level edges (p = 0.054–0.149) with medium effect size (median d ≈ –0.60, range –0.76 to –0.49). These TFNBS findings converge with the cNBS results, demonstrating robust, spatially extensive decreases in gamma FP connectivity after POS ccPAS relative to controls. Together, these gamma band reductions align with our prediction that brain-state-coupled ccPAS leads to decreased FP gamma connectivity.

### Positive ccPAS increased alpha FP connectivity

cNBS (positive tail) revealed significantly greater FP alpha connectivity (Figure 7 E-F) for POS vs mean control (p = 0.015, d ≈ 0.58), with a trend-level increase for POS vs RAND (p = 0.105, d ≈ 0.62). POS vs PREF was above threshold (p = 0.187). TFNBS yielded 26 FWER-significant FP edges for POS vs mean control (p = 0.016–0.044) with small-to-large effect sizes (median d ≈ +0.50, range +0.28 to +0.91), and 10 additional trend-level edges (p = 0.053–0.138) with small-to-medium positive effects (median d ≈ +0.46, range +0.17 to +0.73). POS vs RAND showed 14 trend-level FP increases (p = 0.113–0.13) with medium-to-large effect sizes (median d ≈ +0.86, range +0.62 to +1.19). POS vs PREF was also above threshold (lowest p = 0.187). These alpha band increases are consistent with our prediction of enhanced FP alpha coordination following Positive theta-locked ccPAS.

### No network-level theta modulation

Despite triggering ccPAS on positive theta phases, neither cNBS nor TFNBS detected significant theta FP modulation across contrasts. All p-values exceeded 0.05. These null findings do not support the predicted increase in FP theta connectivity.

### White matter tractography

To disambiguate whether the observed network modulation is supported by direct versus polysynaptic structural pathways, we performed post hoc tractography. To offer an initial anatomical context, we report single-subject tractography, with full group-level quantitative analyses reserved for future work. Figure 8 illustrates a modest direct connection between the stimulated regions via the inferior fronto-occipital fasciculus (IFOF). When either the prefrontal or parietal regions are used as a seed, the tracts terminate at the following common endpoints: inferior frontal gyrus, cerebellum, and anterior/medial temporal areas. This may indicate that stimulation-induced propagation of neural activity is mediated by both direct and indirect axonal connections. Of note, since the mPFC target was functionally identified (subregion with highest theta SNR along the superior frontal gyrus), it is possible that for other subjects a different white matter tract connects both stimulated prefrontal and parietal targets. A candidate alternative tract is the cingulum bundle which also connects subregions of the superior parietal lobule to the medial portion of the superior frontal gyrus (Jitsuishi and Yamaguchi, 2023; Tanglay et al., 2021; Briggs et al., 2020; Wang et al., 2019; Wu et al., 2016; van den Heuvel et al., 2008).

**Figure 8.**
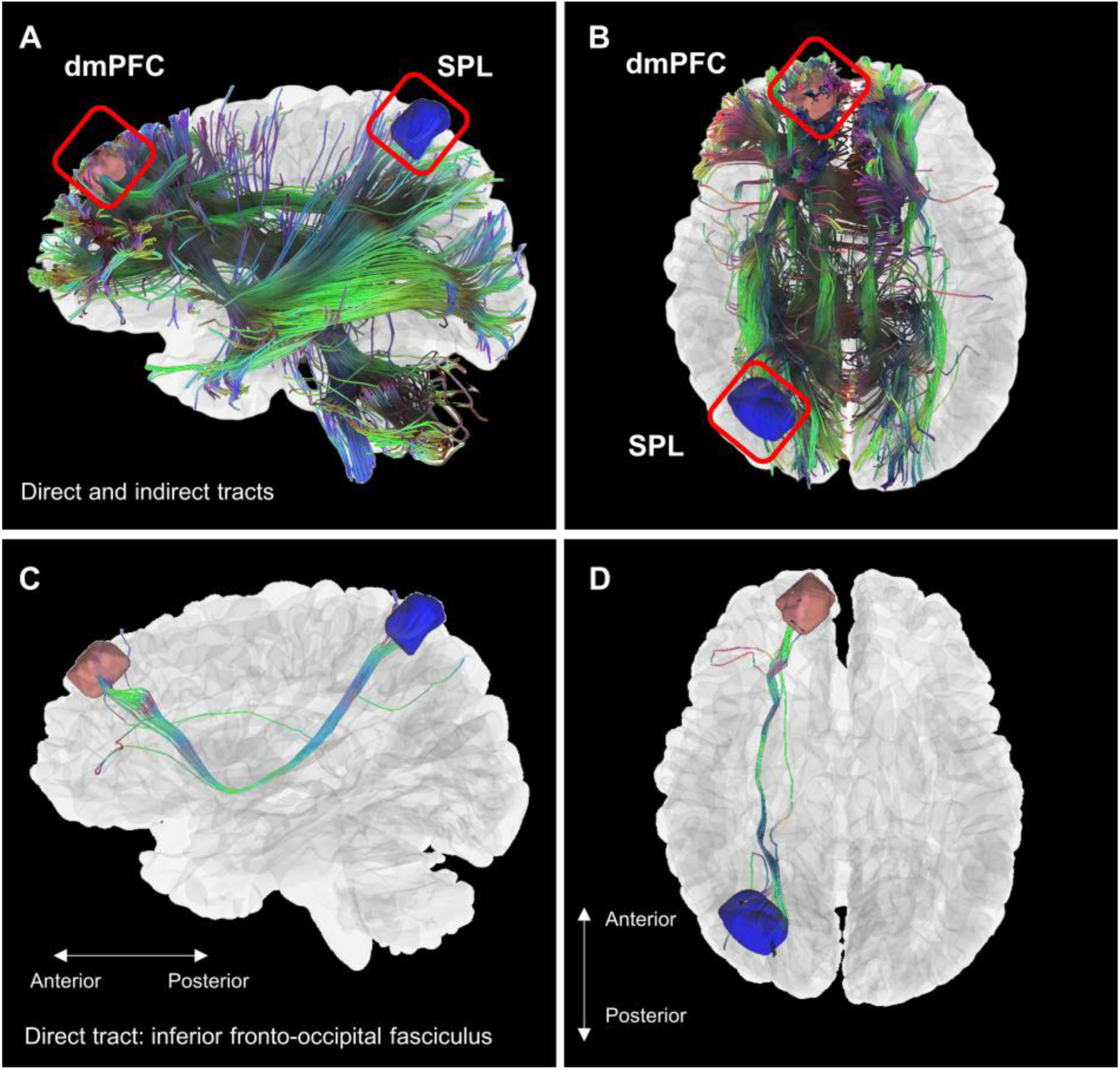
Reconstructed model of the white matter tracts from the stimulated brain regions. White matter tractography from a pre-experiment diffusion weighted MRI scan of a representative subject, with the prefrontal ROI as a seed. Panels A and B show direct and indirect tracts connecting the stimulated prefrontal (red) and parietal (blue) regions. Panels C and D illustrate a modest direct connection via the inferior fronto-occipital fasciculus (IFOF). dmPFC, dorsomedial prefrontal cortex; SPL, superior parietal lobule.

## Discussion

ccPAS is a powerful tool for modulating targeted cortico-cortical pathways; however, extending it to brain-state-coupled delivery entails substantial challenges in targeting accuracy, state estimation, and optimizing dual-site stimulation parameters (Section 1). This paper presents a roadmap with strategies for addressing current challenges and, importantly, introduces a novel application of real-time brain-state-coupled ccPAS in a cognitive network (Section 2). Theta-phase-locked fronto-parietal ccPAS TMS was combined with EEG to test whether delivering stimulation at the positive phase of ongoing mPFC theta would induce distinct changes in evoked EEG activity and FP network connectivity compared to phase-uncoupled ccPAS (RAND) and phase-coupled single-site (PREF) control conditions. At the evoked level, brain-state-coupled ccPAS (POS) produced a fronto-central reversal of the canonical N45 component together with a robust right parieto-temporal N45 negativity relative to both controls. Moreover, functional connectivity measured during the immediate post-intervention period showed that POS ccPAS induced a robust, widespread decrease in gamma FP connectivity and moderate increase in FP alpha connectivity beyond either control ingredient alone. Together, these changes in evoked activity and rapid network reconfiguration may index brain-state-coupled ccPAS efficacy—consistent with “phase-gated STDP”, where oscillatory phase gates cortical excitability and thereby modulates STDPinduction efficacy (Andrade-Talavera et al., 2023; Wang et al., 2023, Roy et al., 2014; Kwag and Paulsen, 2009).

### P30: Early evoked response primarily reflects phase-dependent gain

At P30, only the POS vs RAND contrast showed effects surviving Holm-FWER correction across contrasts x channels x TOIs, localized at prefrontal and posterior electrodes. POS vs PREF and RAND vs PREF showed only uncorrected trends at the same sites. Under the ingredient logic, this pattern is consistent with a phase-dependent *consistency gain*—with stimulation locked to a consistent theta phase, trial-to-trial latency variability decreases; thus, decreasing cancellation during averaging and yielding a larger, more stable averaged TEP. POS and PREF share dmPFC theta phase-locking, whereas RAND disrupts phase alignment. Accordingly, early TEP amplitude/topography can therefore differ between POS and RAND—even if similar generators are recruited under matched stimulation sites and pulse parameters—because phase-locking increases temporal consistency of the evoked response (Zrenner et el., 2018; Desideri et al., 2019).

### N45: Phase-locking plus paired input drive dissociates fronto-central and right parieto-temporal fields consistent with E/I reweighting and stabilized propagation

N45 yielded the most pronounced effects and cleanest dissociation across contrasts. A statistically robust, focal fronto-central reversal of the canonical N45 component emerged when phase-targeting was isolated (POS > RAND), accompanied by a broader, subthreshold fronto-central shift in the same direction when parietal drive was isolated (POS vs PREF). In parallel, right parieto-temporal sites showed a negative deflection during brain-state-coupled ccPAS (POS < RAND, PREF). This is consistent with a redistribution of inhibitory-weighted field contributions across the network—a reduced inhibitory dominance at fronto-central sensors co-occurs with increased or shifted inhibitory dominance in remote parieto-temporal channels. Across topographies, the Holm-surviving versus subthreshold pattern is mechanistically coherent when interpreted in terms of what each contrast isolated. POS vs RAND (parietal drive constant; phase-locking isolated) yields more temporally precise and spatially focal effects with more Holm survivors, whereas POS vs PREF (phase-locking constant; parietal drive isolated) is broader and more variable across subjects, producing many consistent-direction but subthreshold electrodes.

### Phase-locked ccPAS shifts FP gamma and alpha connectivity, not theta

The connectivity analyses demonstrated a striking spectral dissociation. As predicted, POS ccPAS produced robust network-level and edge-level reductions in gamma FP connectivity and increased alpha connectivity relative to control conditions—indicating an FP network state distinct from both controls. The expected increase in theta connectivity, however, was not observed. These frequency-specific reconfigurations in resting state connectivity following POS ccPAS suggest that the post-intervention network state reflects the interaction of phase-alignment with associative pairing, rather than either ingredient alone. The divergent changes in alpha and gamma synchrony align with prior work demonstrating that these rhythms engage feedback and feedforward processing, respectively (Michalareas et al., 2016; Fries, 2015; van Kerkoerle et al. 2014). The absence of theta FP modulation may seem surprising given that stimulation was locked to prefrontal theta phase. A plausible interpretation is that theta functions as a control variable that gates excitability and spike timing during stimulation, with the durable network reconfiguration primarily expressed in alpha and gamma bands—consistent with cross-frequency mechanisms proposed for working memory and cognitive control (Palva et al., 2010; Siebenhühner et al., 2016; Fernández et al., 2021; Martinez-Cancino et al., 2019).

### Phase-gated associative plasticity induction: Mechanistic and cognitive implications

The POS ccPAS protocol was designed to instantiate associative, STDP-like modulation by combining a fixed inter-pulse interval with brain-state targeting intended to increase effective temporal precision. First, the 10 ms ISI between parietal and prefrontal pulses align with human ccPAS evidence that pathway modulation depends on pulse order and timing, consistent with STDP-like rules (Nord et al., 2019; Momi et al., 2019; Santarnecchi et al., 2018; Casula et al., 2016; Koch et al., 2013). Second, in our ingredient-control framework, POS vs RAND isolates phase-locking while POS vs PREF isolates dual-site associative drive. Correspondingly, the predominance of post-intervention connectivity reconfiguration in POS relative to both controls supports input- and timing-contingent specificity—arguing against a generic dmPFC excitability shift mechanism. Third, critically, these fronto-parietal connectivity reconfigurations were expressed during a 7-minute post-intervention resting-state recording, supporting initial persistence— short-term plasticity expression beyond the induction period. Short-term plasticity is known to alter routing and effective strength within neural circuits, thereby dynamically reconfiguring functional connectivity on timescales of milliseconds to minutes (Zucker and Regehr, 2002; Abbott and Regehr, 2004; Citri and Malenka, 2008). Future work will distinguish induction and early expression from consolidation by tracking whether the early network reconfiguration persists, decays, or strengthens at later time points, consistent with longer-term potentiation.

Mechanistically, the findings are interpreted as evidence for phase-gated STDP induction with short-term network-level expression, rather than durable LTP. Within this framework, oscillatory phase biases excitability and responsiveness, thus, increasing the reliability and potency of STDP-like pairing in modulating the targeted pathway. Theta phase-dependent gating of hippocampal plasticity is well-established, with stimulation delivered at different phases of the theta cycle producing LTP or LTD (Schall et al., 2008; Hyman et al., 2003; Huerta and Lisman, 1995; Hölscher et al., 1997). Moreover, prefrontal neurons can phase-lock to hippocampal theta (Siapas et al., 2005), providing support that theta phase can coordinate prefrontal network responsiveness. Consistent with this, real-time EEG-informed stimulation studies show that targeting specific oscillatory phases can modulate responsiveness and plasticity-relevant outcomes (Zrenner et al., 2018; Stefanou et al., 2018). Notably, previous work in computational models and hippocampal slices have demonstrated that the timing of an external input relative to the oscillatory phase can bidirectional gate STDP direction and behavior. (Wang et al., 2023; Andrade-Talavera et al., 2023, Roy et al., 2014; Kwag and Paulsen, 2009), providing computational and in-vitro evidence of phase-gated STDP. While STDP-like plasticity has been demonstrated in humans and oscillatory phase has been shown to gate cortical excitability, no prior study has provided direct systems-level experimental evidence consistent with phase-gated STDP in humans. The present findings bridge these literatures by showing that associative plasticity induced by ccPAS is modulated by theta oscillation phase, consistent with oscillatory gating of STDP induction.

In this context, the POS-associated increase in trial-to-trial stability at P30 and the fronto-central N45 polarity reversal provide convergent physiological signatures of successful phase-locked associative engagement during stimulation. Pharmacological TMS-EEG studies indicate that N45 is sensitive to GABAergic and glutamatergic mechanisms. (Premoli et la. 2014; Belardinelli et al., 2021); thus, we propose that the N45 polarity reversal reflects a reweighting of E/I balance away from inhibitory dominance and toward facilitation—positioning it as a candidate biomarker of phase-locked associative plasticity induction. The subsequent frequency-specific changes in FP connectivity is compatible with a rapid reweighting of large-scale communication and gating, given the roles of gamma and alpha synchrony in mediating local and inter-regional coordination and inhibition (Fries, 2015; Jensen & Mazaheri, 2010; Palva & Palva, 2011).

Together, these evoked and post-intervention network effects support a constrained mechanistic interpretation in which theta-phase targeting stabilizes the timing of local responsiveness and its propagation, thereby enabling STDP-like pairing rules to more consistently bias connectivity within the FP architecture. The FP network is a core hub for cognitive control and working memory, integrating sensory inputs with executive computations (Palva et al., 2010; Langer et al., 2013; Phillips et al., 2014; Jacob et al., 2018; Goddard et al., 2022). We propose that phase-gated STDP—a distinct form of associative plasticity—provides a plausible pathway by which phase-locked ccPAS can influence cognitive control circuity, bridging synaptic timing rules with changes in large-scale FP network architecture which has been shown to shape subsequent task-evoked reconfiguration (Cole et al., 2013; Bassett et al., 2011).

### Methodological considerations and limitations

A number of methodological issues qualify our conclusions. The first is the limited sample size and that the PREF condition was added post-hoc at the reviewers’ request which may limit power for some contrasts. Nonetheless, the gamma connectivity decreases vs both PREF and mean control were large and FWER-significant at edge level, and the alpha connectivity increases were robust at the family level. The use of permutation-based cNBS/TFNBS with explicit baseline comparability checks help mitigate concerns about inflated false positives (Noble and Scheinost, 2020; Baggio et al., 2018; Zalesky et al., 2010; Maris & Oostenveld, 2007). We are also addressing this in an expanded study involving 20 subjects with additional experimental conditions and evaluating stimulation-induced effects on working memory (Jovellar et al., in preparation). Second, wPLI as a connectivity measure cannot determine directionality of information flow. Directed measures and combined TMS-EEG-fMRI (Peters et al., 2020) approaches could help disambiguate whether prefrontal-to-parietal interactions or vice versa drive the observed alpha/gamma changes. Third, assessing the anatomical connections between the stimulated prefrontal and parietal regions was done post-hoc. Future experiments could perform white matter tractography in real-time during the stimulation session (Aydogan et al., 2025) or offline before the start of the ccPAS interventions to improve anatomical precision.

### Future directions

Near-term work will prioritize validating the N45 polarity reversal as a biomarker of phase-locked associative plasticity induction by: *(i)* conducting source-reconstructed ccPAS-TEP analysis; *(ii)* examining whether online ccPAS-evoked responses generalize to offline single-pulse TEP (mPFC, SPL); and *(iii)* testing whether individual differences in the online N45 signature predict post-intervention network reconfiguration. In parallel, causal inference for a phase-gated STDP mechanism can be strengthened by: *(i)* performing directed connectivity analyses; *(ii)* assessing long-term plasticity by tracking connectivity modulation at later time points and testing for concomitant behavioral effects, and *(iii)* (where feasible) pharmaco–TMS–EEG validation to test whether POS selectively modulates directed fronto-parietal influences and whether N45/connectivity effects are attenuated under GABA and NMDA manipulations. Mid-term, state-coupled targeting should expand beyond phase-based triggering to multi-feature state targeting and explore other closed-loop control techniques. Finally, next-generation implementations include: *(i)* multi-locus TMS (mTMS) which enable rapid multi-site stimulation where stimulation parameters can be adjusted at a millisecond scale (Navarro de Lara et al., 2021; Koponen et al., 2018; Nieminen et al., 2022; Souza et al., 2022); and *ii)* adaptive closed-loop brain stimulation that allows precise fine-tuning by flexibly adapting stimulus parameters in response to dynamic shifts in brain activity (Humaidan et al., 2024; Sherf et al., 2021) that may occur during the stimulation session.

## Conclusions

In summary, this paper presents the first implementation of brain-state-coupled ccPAS in a cognitive network and includes a methodological roadmap for improving targeting accuracy and ease of application. Beyond serving an illustrative protocol, this work introduces and provides empirical evidence for the induction and early expression of phase-gated STDP in humans. Our work provides a crucial first step toward personalized pathway-targeted brain stimulation in cognitive networks, potentially enhancing circuit specificity while minimizing off-target effects. This framework offers a foundation for developing individualized interventions in people with network impairments affecting memory, such as Alzheimer’s disease, major depression, or ADHD, as well as optimizing cognitive enhancement approaches in healthy individuals.

## Conflict of interest statement

The authors declare no competing financial interests.

## Acknowledgments

DBJ was awarded the Carl Duisberg Fellowship for Medical Sciences in support of the work reported in this manuscript. This work was part of the ConnectToBrain consortium and was supported by the European Research Council Synergy Grant 810377 awarded to UZ and collaborators. Funders had no role in the preparation of this manuscript or the decision to publish.

## Author contributions

DBJ: Conceptualized the research design, wrote the experiment and analysis codes, performed experiments, analyzed data, wrote the initial and final drafts of the manuscript. ST: Wrote the initial drafts of the manuscript, reviewed and edited the manuscript. PB: Provided tools for source localization, reviewed and edited the manuscript. OR: White matter tractography, reviewed and edited the manuscript. ES: Reviewed and edited the manuscript. UZ: Refined the research design, acquired funding, reviewed and edited the manuscript.

## Data availability

Analysis codes are available upon request to the corresponding author. Public sharing of raw data is not possible due to the data protection agreement with the participants. Pre-processed data are available upon request to the corresponding author.

## Notes

### Competing Interest Statement

The authors have declared no competing interest.

