## Supplemental Information for "Real-time brain-state-coupled cortico-cortical paired associative stimulation of cognitive networks"

### SUPPLEMENTAL RESULTS

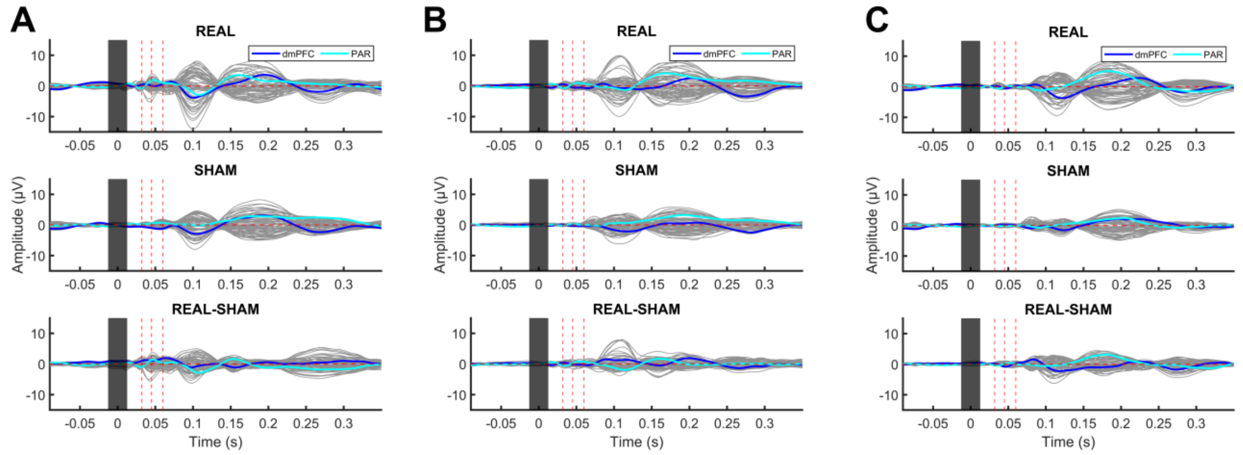

**Figure S1 | TMS-evoked potentials for POS, RAND, and PREF (0–350 ms).**
(A–C) Butterfly plots of TMS-evoked potentials (µV) are shown for POS (A), RAND (B), and PREF (C), separately for REAL (top), SHAM (middle), and REAL–SHAM (bottom; sham template subtraction). Gray traces show channel-wise waveforms. Colored traces show ROI time courses for electrodes surrounding the stimulated sites: dmPFC (blue; Fz, F1, F2, AFz, AF3, AF4) and PAR (cyan; CP1, CPz, CP2, P1, Pz, P2). The dark bar marks the TMS pulse/artifact window, and dashed red vertical lines denote the analyzed time windows of interest (TOIs: 30, 45, and 60 ms). This figure is descriptive; inferential statistical evaluation is reported in the main Results figures. *dmPFC*, dorsomedial prefrontal cortex; *PAR*, parietal; *TOI,* time window of interest; *s*, seconds; *POS*, Positive ccPAS; *RAND*, Random ccPAS; *PREF*, Prefrontal only TMS

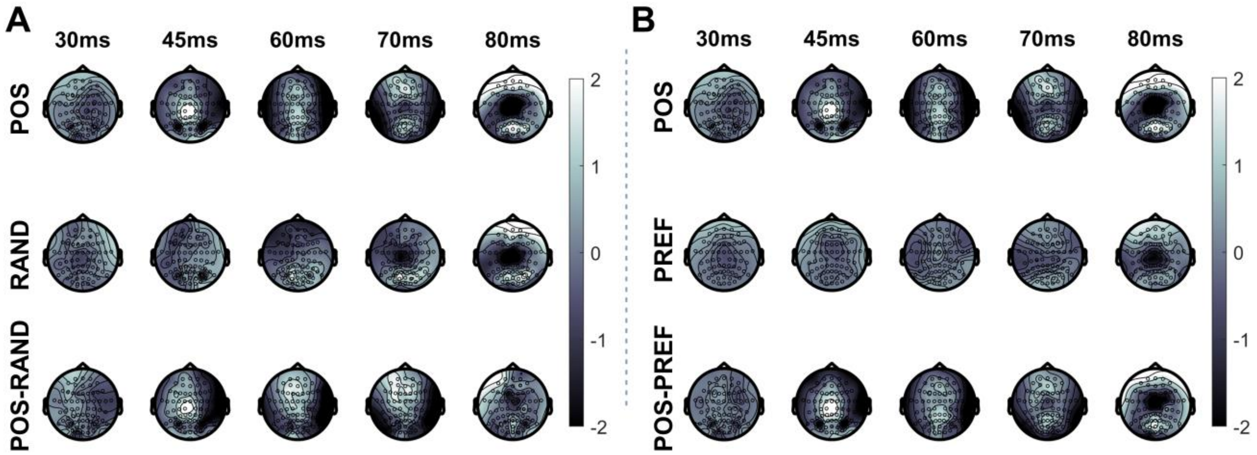

**Figure S2 | Real-only (non–sham-subtracted) scalp topographies and condition differences.**
(A) Grand-average Real-only TMS-evoked scalp potentials (µV) are shown for POS (top row), RAND (middle row), and their difference (POS − RAND; bottom row).
(B) Grand-average Real-only TMS-evoked scalp potentials (µV) are shown for POS (top row), PREF (middle row), and their difference (POS − PREF; bottom row).
Columns correspond to 30, 45, 60, 70, and 80 ms after the TMS pulse. Color scales are shown to the right of each panel. These maps provide a descriptive visualization of the spatial distribution of evoked activity and condition differences before sham-template subtraction; inferential statistical analyses were performed on the cleaned REAL–SHAM signal and are reported elsewhere.*POS*, Positive ccPAS; *RAND*, Random ccPAS; *PREF*, Prefrontal only TMS

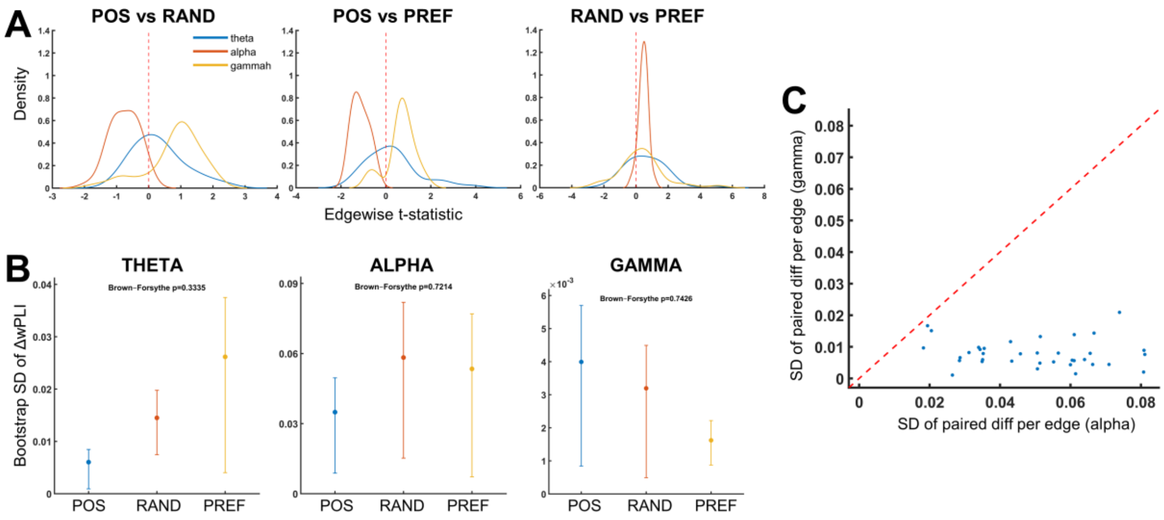

**Figure S3 | Baseline fronto-parietal comparability and Δ-variance structure support exchangeability for permutation-based connectivity inference.**
(A) Baseline (PRE) FP edgewise comparability diagnostics. Kernel density estimates of edgewise t-statistics for FP edges are shown for POS vs. RAND (left), POS vs. PREF (middle), and RAND vs. PREF (right), overlaid by band [theta (4–8 Hz), alpha (9–12 Hz), and gamma (60–90 Hz)]. Distributions are centered near zero, indicating no large systematic baseline bias across contrasts. The red dashed line marks t = 0.
(B) Bandwise variance diagnostics for subject-level FP mean ΔwPLI (POST–PRE). For each condition, the SD of subject-level FP mean ΔwPLI was estimated by bootstrap resampling subjects with replacement (n = 2000); points denote the median bootstrap SD and whiskers denote the 2.5th and 97.5th percentiles (95% bootstrap interval). Brown–Forsythe tests indicated no evidence of unequal variance across POS, RAND, and PREF in theta (p = 0.3335), alpha (p = 0.7214), or gamma (p = 0.7426).
(C) Edgewise Δ-variance structure across FP edges. The SD across paired subjects of the edgewise paired Δ differences (POS–RAND) is plotted for alpha (x-axis) versus gamma (y-axis). Each point corresponds to one FP edge, and the red dashed diagonal indicates the identity line. Most points fall below the identity line, indicating that alpha edges generally showed greater Δ variability than gamma edges, without evidence of extreme or irregular heterogeneity. Together, these diagnostics support the exchangeability assumptions required for permutation-based cNBS and TFNBS. *FP*, fronto-parietal; *dwPLI*, debiased weighted phase lag index; *cNBS,* constrained network-based statistic; *TFNBS*, threshold-free network-based statistics; *POS*, Positive ccPAS; *RAND*, Random ccPAS; *PREF*, Prefrontal-only TMS.

| Frequency | Contrast | Observed stat | p_value | d | CI low | CI high |
|---|---|---|---|---|---|---|
| Gamma | POS_vs_MEANCTRL | 1.101307554 | 0.011899 | -0.87334 | -2.50416 | -0.51477 |
| | POS_minus_RAND | 1.152437647 | 0.041796 | -0.68145 | -0.92089 | -0.03971 |
| | POS_minus_PREF | 0.874201638 | 0.048795 | -1.15682 | -2.31636 | 0.119374 |
| Alpha | POS_vs_MEANCTRL | 1.47159989 | 0.025997 | 0.495707 | -0.2288 | 1.464794 |
| | POS_minus_RAND | 1.034647504 | 0.113689 | 0.642447 | -0.51541 | 3.176189 |
| | POS_minus_PREF | 0.726613058 | 0.266573 | 0.548899 | -0.69108 | 1.463368 |

**Table S1 | Secondary ROI cNBS analysis preserves frequency-specific fronto-parietal reconfiguration after brain-state-coupled ccPAS: ↓ gamma, ↑ alpha connectivity.** Family-constrained cNBS results are shown for a secondary fronto-parietal ROI set used to assess the robustness of the main connectivity findings for gamma and alpha. Rows correspond to contrasts [POS vs. mean control (average of PREF and RAND), POS vs. PREF, and POS vs. RAND]. Reported values include the observed family statistic, permutation p value, Cohen's d, and 95% confidence interval. *Top:* In the gamma band, cNBS indicates reduced family-level connectivity after POS ccPAS for POS vs. mean control ($p = 0.0119$, $d \approx -0.87$), POS vs. RAND ($p = 0.0418$, $d \approx -0.68$), and POS vs. PREF ($p = 0.0488$, $d \approx -1.16$). *Bottom*: In the alpha band, cNBS indicates increased family-level connectivity after POS ccPAS for POS vs. mean control ($p = 0.0260$, $d \approx 0.50$]), no effect for POS vs. PREF ($p = 0.2666$, $d \approx 0.55$), and a trend-level effect for POS vs. RAND ($p = 0.1137$, $d \approx 0.64$). Overall, the secondary ROI-set analysis preserved the main spectral pattern observed in the primary analysis, with reduced gamma connectivity across contrasts and stronger alpha increases for POS vs. mean control than for the single-control contrasts. cNBS provides family-wise-error-rate-controlled inference at the FP-family level. MEANCNTRL, mean control; cNBS, constrained network-based statistic.

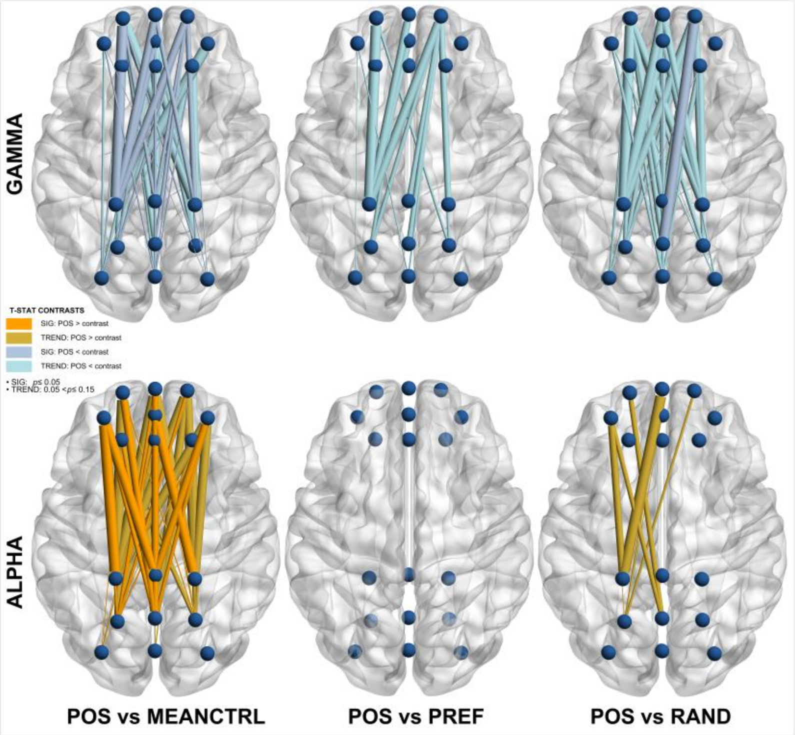

**Figure S4 | Frequency-specific fronto-parietal reconfiguration is consistent across ROI definitions.** TFNBS edgewise connectivity diagrams are shown for a secondary predefined fronto-parietal ROI set used to assess the robustness of the main connectivity findings. Rows correspond to frequency band [top, gamma (60–90 Hz); bottom, alpha (9–12 Hz)], and columns correspond to contrast [left, POS vs. mean control (average of PREF and RAND); middle, POS vs. PREF; right, POS vs. RAND]. Edge colors follow the same conventions as in Figure 7: darker warm colors indicate significant increases after POS ($p_{FWER} \leq 0.05$), darker cool colors indicate significant decreases after POS ($p_{FWER} \leq 0.05$), and lighter shades indicate trend-level effects ($0.05 < p_{FWER} < 0.15$). *Top:* Gamma band TFNBS edgewise connectivity diagrams show reduced connectivity after POS ccPAS. POS vs. mean control shows 23 significant ($p = 0.018–0.037$) and 21 trend-level edges ($p = 0.053–0.142$). POS vs. PREF shows no FWER-significant edges and 14 trend-level edges ($p = 0.053–0.139$). POS vs. RAND shows 3 significant ($p = 0.048$) and 35 trend-level edges (p = 0.055–0.150). *Bottom:* Alpha band TFNBS edgewise connectivity diagrams show increased connectivity after POS ccPAS. POS vs. mean control shows 21 significant (p = 0.028–0.044) and 28 trend-level edges (p = 0.052–0.138). POS vs. PREF shows no edges below threshold. POS vs. RAND shows 8 trend-level edges (p = 0.122–0.129). Overall, the secondary ROI-set analysis preserved the main spectral pattern observed in the primary analysis. Nodes correspond to electrodes in the secondary predefined frontal and parietal ROI set. TFNBS provides family-wise-error-rate-controlled edgewise inference. MEANCNTRL, mean control; TFNBS, threshold-free network-based statistics.

### SUPPLEMENTAL METHODS

#### Text S1 | Diffusion MRI processing and tractography

The diffusion-MRI processing pipeline used to reconstruct white-matter pathways linking the individualized dmPFC and SPL targets are as follows. T1-weighted images were segmented and meshed (FieldTrip) and used to build EEG boundary-element head models (Oostenveld et al., 2011; Stenroos & Sarvas, 2012; Stenroos & Nummenmaa, 2016). Diffusion MRI (3T Siemens Prisma; multi-shell b = 900/1600/2500 s/mm²; ~18/32/50 directions; 1.5 mm isotropic) underwent susceptibility-distortion correction with FSL TOPUP, then reconstruction and tractography in DSI Studio using generalized q-sampling imaging (GQI) and deterministic streamline tracking with quantitative anisotropy (QA) (Yeh et al., 2010; Yeh et al., 2013). Seed ROIs were defined in native (subject) T1 RAS space as spheroids centered on the subject-specific dmPFC and SPL stimulation targets. We used high-density seeding, step size = voxel, curvature threshold 45–90°, and retained streamlines 30–300 mm. Topology-informed pruning (TIP) was applied to reduce likely false positives (Yeh et al., 2019). These reconstructions are illustrative—intended to provide anatomical context for potential pathways rather than formal connectomic inference.

#### Text S2 | TMS-evoked potential statistics

Statistical analyses of ccPAS-TEP/TEP data used a partially paired general linear model (GLM) with permutation-based inference. The model was fit by ordinary least squares (OLS) and included subject indicator regressors for participants contributing observations to both conditions (repeated IDs). To compare conditions under a partially paired/partially overlapping design (no duplicate observations within a condition; possible subject overlap across conditions), we used nonparametric permutation inference implemented in FieldTrip (`ft_timelockstatistics`, `cfg.method='montecarlo'`) with a patched Monte Carlo engine using a custom permutation schedule (cfg.resample) that preserves exchangeability under partial overlap. This inferential setting corresponds to the partially paired/partially overlapping mean-comparison problem reviewed in the methodological literature (Guo and Yuan, 2017; Derrick et al., 2017), which also includes likelihood-based approaches such as the modified MLE of Lin and Stivers (1974).

For each analysis, we constructed a two-row FieldTrip design matrix with condition labels (`design(1,:)`) and subject identifiers (`design(2,:)=subject_num`). For the three-group omnibus (A/B/C), the test statistic was an *F-statistic* from a reduced-vs-full OLS GLM (`statfun_glm_partial`) in which the reduced model included the intercept and repeated-ID subject regressors, and the full model additionally included condition regressors for the non-reference groups; degrees of freedom were computed from the model ranks (`df1 = rank(X_F) − rank(X_R)`, `df2 = Nobs − rank(X_F)`). For two-condition contrasts (A vs B), the test statistic was the *t-value* of the condition coefficient from an OLS GLM (`statfun_glm_partial_t`) including an intercept, subject indicator regressors for repeated IDs only, and a binary condition regressor (e.g., `Xc = double(cond==1)`), with degrees of freedom `df = Nobs − rank(X)`.

Inference used the exchangeability-constrained permutation procedure matched to the partial-overlap structure. A permutation plan `resample` (`K=nperm`, fixed seed) was generated per contrast/TOI and applied in FieldTrip by reordering the design columns at each randomization (`tmpdesign = design(:, resample(i,:))`) while leaving the data matrix unchanged in the permutation branch. The plan (i) performs within-subject swaps/permutations for repeated-ID blocks and (ii) permutes singleton observations among singletons only. Two-condition contrasts used two-sided p-values (`cfg.tail=0`, `cfg.correcttail='prob'`), whereas omnibus F-tests used a right-tailed test (`cfg.tail=1`; `cfg.correcttail='prob'`).

Multiple-comparison control used the uncorrected Monte Carlo p-values (`cfg.correctm='no'`) followed by manual correction. Within each TOI, p-values were adjusted across the within-family set of tests (channels for time-averaged analyses; channels×time for channel-by-time analyses) using Holm correction. For the two-condition pipeline, we additionally controlled global FWER by pooling uncorrected p-values across channels × TOIs × contrasts and applying Holm correction to the pooled family, yielding `p_global`; channels were declared significant at `p_global ≤ α` (α=0.05). For descriptive reporting, we extracted condition-wise means and differences from TOI-averaged data. For two-condition contrasts we computed signed r-equivalent effect sizes from t and df (`r = sign(t)*sqrt(t^2/(t^2+df))`), and for omnibus tests we computed partial eta-squared from F and its degrees of freedom (`η²p = (df1·F)/(df1·F + df2)`).

#### Text S3 | Functional connectivity quantification and statistics

The functional connectivity analyses were conducted using custom MATLAB scripts implementing the pipeline described below:

*Resting-state connectivity matrices and ROI graph*

Resting-state EEG trials were separated into PRE and POST epochs and were downsampled to 500 Hz prior to spectral estimation. Band-limited Fourier spectra were then computed in FieldTrip (Oostenveld et al., 2011) using a multi-taper approach across theta (4–8 Hz), alpha (8–13 Hz), and gamma (60–90 Hz) bands. Connectivity was estimated from the Fourier spectra with the debiased weighted phase lag index and then averaged across trial/taper repetitions to obtain one PRE and one POST connectivity matrix per subject and condition. This choice follows the use of multitaper spectral estimation for electrophysiological data and of debiased wPLI to reduce sensitivity to near-zero-lag contamination and sample-size bias (Mitra and Pesaran, Biophysical Journal, 1999; Vinck et al., NeuroImage, 2011; Oostenveld et al., Computational Intelligence and Neuroscience, 2011). ([PMC][1])

The primary confirmatory ROI graph comprised frontal nodes `Fp1`, `Fp2`, `Fpz`, `AFz`, `AF3`, `AF4` and parietal nodes `CPz`, `CP1`, `CP2`, `Pz`, `P1`, `P2`. A larger ROI graph (`Fp1`, `Fp2`, `Fpz`, `AFz`, `AF3`, `AF4`, `Fz`, `F1`, `F2` with `CP1`, `CP2`, `CPz`, `Pz`, `P1`, `P2`, `POz`, `PO3`, `PO4`) was reserved for supplemental robustness analyses. Subject-level change matrices were defined as Δ = POST − PRE and vectorized as the upper triangle of the ROI graph, which naturally partitioned edges into frontal-frontal (FF), parietal-parietal (PP), and fronto-parietal (FP) families. The FP family was used downstream.

*Contrast-specific edgewise test statistics and permutation maps*

Observed edgewise statistics were computed from subject-level change scores (Δ = POST − PRE) for the planned contrasts of Positive versus mean control, POS vs. RAND, and POS vs. PREF. For each contrast, edgewise subject matrices were assembled by condition and aligned by subject identifier so that paired and partially overlapping observations could be accommodated explicitly at the edge level. The POS vs. RAND and POS vs. PREF pairwise contrasts were both analyzed within the same partially overlapping-samples framework, using row-stacked two-condition data and an overlap-aware studentized statistic with Welch-type variance handling, raw overlap-based covariance scaling, and PSD-based variance repair. For the POS vs. mean-control contrast, the observed edgewise map was computed with an overlap-aware three-condition studentized contrast targeting Δ_POS − 0.5(Δ_OPEN + Δ_PREF). This procedure incorporated a minimum-overlap rule for covariance estimation, a small variance floor for numerical stability, and PSD-based variance repair. The partially overlapping statistics are directly grounded in the Derrick partially overlapping-samples framework, which interpolates between paired and independent-samples t procedures while retaining all available data (Derrick et al., 2017).

Permutation maps were generated using contrast-matched null procedures with 10,000 permutations, a fixed random seed, and one-sided tail inference. For the POS vs. mean-control contrast, condition labels were permuted at the row level within the full POS/RAND/PREF row universe while preserving within-subject exchangeability and, where applicable, shuffling singleton labels only across singleton rows. The POS vs. RAND and POS vs. PREF pairwise contrasts were both evaluated within the same partially overlapping permutation framework, with permutations restricted to the corresponding two-condition row universe. This stage yielded, for each contrast, an observed edgewise statistic vector and a matched permutation ensemble defined on the same ROI edge universe.

*Network-level inference*

<u>Family-constrained cNBS analysis</u>

A family-constrained network statistic adapted from the cNBS framework was used to test whether the Positive condition produced distributed effects across the FP edge family (Noble and Scheinost, 2020). After applying the same tail transformation used for the edgewise maps, the transformed edgewise t statistics were aggregated within the FP family using their mean. Using the contrast-specific permutation maps described above, the observed FP family statistic was compared with the corresponding null distribution of permuted FP family statistics. Because the confirmatory analysis was restricted to this single predefined family, the reported p value is a single-family permutation p value.

<u>Threshold-free network-based statistics (TFNBS)</u>

In parallel, TFNBS was applied within the selected FP edge subset using an edge-adjacency graph in which two edges were defined as neighbors if they shared a node. Following the TFNBS logic, edgewise statistic maps were tail-transformed and enhanced across thresholds without a fixed component-defining threshold (Smith and Nichols, 2009; Baggio et al., 2018). Enhancement was accumulated as $\text{extent}^E \times \text{tau}^H \times d\tau$, with extent defined by suprathreshold connected-component size ($E = 0.9$, $H = 0.5$, fixed threshold grid, $dh = 0.1$). Using the contrast-specific permutation maps described above, a max-null distribution was constructed by recomputing the enhanced map for each permutation and retaining the maximum enhanced value across edges. Observed enhanced edge statistics were then compared with this max-null distribution to obtain one family-wise-error-corrected p value per edge. Network diagrams were visualized using BrainNet Viewer (Xia et al., 2013).

*Effect sizes*

Descriptive effect sizes were computed after inferential testing and were not used to define significance. For the family-constrained analysis, family-level effect sizes were computed from subject-wise mean Δ values averaged across all edges in the family and matched to the estimand of the corresponding contrast. For TFNBS, edge-level Cohen-type effect sizes were estimated for all edges. For the two-condition contrasts, these effect sizes were derived from the subject-level Δ distributions contributing to each edge; for the POS vs. PREF contrast, paired-overlap `dz` values were additionally retained for the intersected subjects. For the POS versus mean control contrast, the primary edge effect size was defined to match the condition-weighted estimand, Positive minus the average of RAND and PREF, and an auxiliary `dz` was computed only for subjects contributing data to all three conditions. Confidence intervals were estimated by bootstrap resampling (2,000 draws; 95% confidence intervals). In addition to the confirmatory threshold of $\alpha = 0.05$, the TFNBS pipeline retained descriptive trend-level results for edges with `0.05 < p_FWER ≤ 0.15`.
